# A Gene-Program Architecture of Mouse T cells

**DOI:** 10.64898/2026.09.15.751918

**Authors:** Ziang Zhang, Tianze Wang, Siba Smarak Panigrahi, Peter Carbonetto, Matthew Stephens, Christophe Benoist, Sara Mostafavi, Maria Brbić, David Zemmour, the immgenT Project

## Abstract

We present a gene-program framework for resolving mouse T cell heterogeneity across the immgenT atlas. On ∼800,000 T cells spanning lineages, organs, and immune challenges, we defined 200 reproducible gene programs, discovered by empirical Bayes matrix factorization approach and validated through a new deep learning approach, that capture major axes of T cell variation, including lineage identity, activation states and tissue location. Gene-program analysis complemented cluster-based annotation by decomposing T cell states into “molecular modules”, revealing quantitative, shared, modules not represented with discrete labels alone. Across tissues, gene programs reflected both tissue-imposed programs and changes in cluster composition. Integrating GP activity with cell-surface marker expression from the CITE-seq data, revealed that markers can report different programs depending on lineage and context. Together, immgenT-GP extends the atlas from a map of T cell states to a molecular reference of the programs that underlie them.

## Introduction

The identification and characterization of coordinated gene-expression into programs (GP) or modules have been central to the interpretation of genome-wide transcription, in immunology and other fields^1–3^. Early studies, which relied on bulk transcriptomic data, inferred gene co-variation across datasets containing diverse cell types. The advent of single-cell technologies has enabled the identification of coordinated gene expression from cell-to-cell variation within a single cell-type, providing a more refined view of cellular states and regulatory programs^4,5^. These efforts aim to link co-expressed genes, that may have diverse molecular roles, e.g., transcription factors, enzymes, or secreted molecules, to integrated cellular functions such as tissue localization, cytokine production or proliferation. They also seek to define the regulation of these functions by transcription factors^6^. This understanding has also informed immunoengineering approaches, including chimeric antigen receptor (CAR) T cells^7^, gene therapy^8^ and T cell reprogramming^6^.

The Immunological Genome (ImmGen) Project established comprehensive transcriptional and chromatin maps across immune cell types, laying the groundwork for understanding immune regulatory networks^9–11^. The immgenT project^12^ has expanded this effort by profiling T cells across hundreds of samples and hundreds of thousands of single cells spanning tissues, developmental stages, infections, tumors, autoimmune settings, and homeostasis. The resulting saturation of T cell states across biological contexts creates a unique opportunity to systematically analyze shared and context-specific gene programs. In the integrated immgenT atlas, cells are mapped into a latent space that captures molecular similarity across experiments and perturbations^12^. Although these latent representations effectively organize cells across biological contexts, they are not directly biologically interpretable. Here, we sought to derive alternative representations that resolve the gene programs underlying T cell identities, states, and phenotypes. Such a GP framework also complements cluster-based analysis by capturing biological variation that is not well represented by discrete labels, including transcriptional programs shared across otherwise distinct clusters and programs that vary among cells within the same cluster.

Although the ultimate objective is to decode the gene regulatory networks^13,14^ (GRNs) and transcription factor logic governing T cell differentiation, a more tractable objective in large-scale single-cell datasets is to identify recurrent patterns of gene co-expression. These patterns, variably referred to as gene modules, gene programs, or transcriptional programs, can be inferred using linear approaches such as non-negative matrix factorization (NMF)^15–21^, and, more recently, non-linear, AI-based models^22,23^. Among these approaches, sparse MF-based methods have been particularly useful in single-cell biology, e.g., scNMF^5,21^, scHPF^17^, because they yield relatively interpretable programs. Empirical Bayes Matrix Factorization (EBMF) and its recent implementation in *flashier*^*24,25*^ extend and generalize these approaches by operating effectively on log-normalized data, accounting for sequencing depth, and allowing flexible priors that encompass NMF, semi-NMF, and related factorization frameworks.

Here, as a complementary approach to the cell-state analyses presented in the companion immgenT manuscripts^12,26–29^, we used EBMF to identify gene programs across the atlas. These programs revealed a modular architecture of T cell states, with transcriptional programs shared across lineages, tissues, and activation contexts. We further validated this structure using Residual Quantized Variational Inference (RQVI), a new deep-learning algorithm for gene-program discovery, and linked program activity to surface-protein expression through paired CITE-seq measurements.

## Results

### Sparse and reproducible gene programs across the immgenT atlas

To capture the breadth of T cell heterogeneity, the immgenT project profiled T cells from 734 samples and ∼800,000 cells, measuring expression of over 20,000 RNA-coding genes and 128 surface proteins^12^. Integration using totalVI^30^, a deep learning framework that jointly models RNA and protein measurements, comprehensively captured T cell heterogeneity (see companion manuscripts^12,26–29^), delineating eight major T cell lineages: CD8, CD4, Treg, gdT cells, CD8aa T cells, Zbtb16+ unconventional T cells (Tz), double-negative (DN) and double-positive (DP) T cells. Each lineage contained 6–21 robust clusters that were reproducible in the RNA, protein, and sample spaces (**Fig. 1a**).

**Figure 1.**
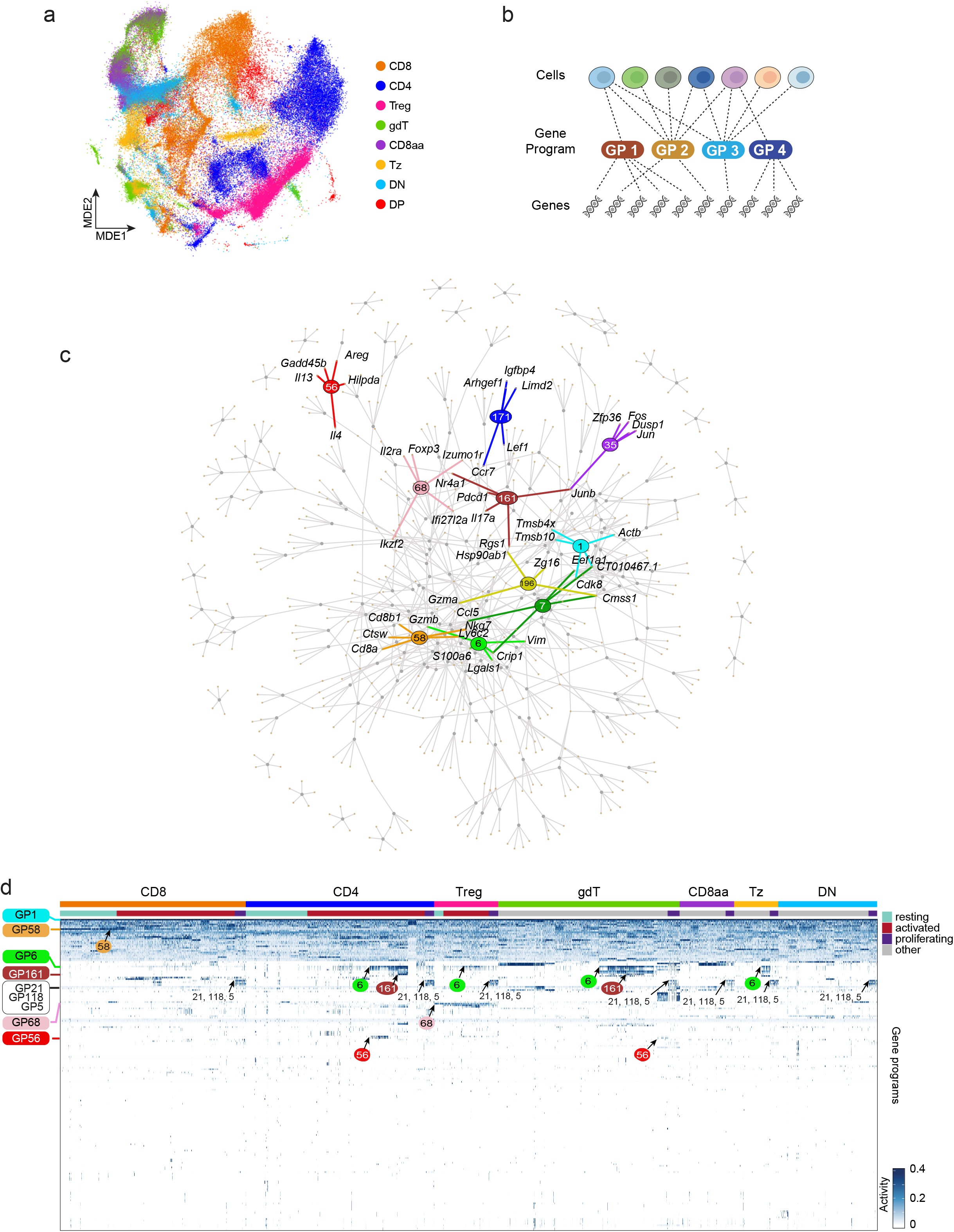
A gene-program architecture underlies T cell heterogeneity across the immgenT atlas. **(a)** Minimum distortion embedding (MDE) plot of all mature T cells profiled in immgenT (all-T MDE), colored by T cell lineage: CD4, conventional CD4^+^ T cells; CD8, conventional CD8αβ^+^ T cells; Treg, CD4^+^ Foxp3^+^ regulatory T cells; CD8aa, CD8aa αβ T cells; gdT, gdT cells; Tz, Zbtb16^+^ αβ T cells; DN, CD4^−^CD8αβ^−^ αβ T cells; DP, CD4^+^CD8αβ^+^ αβ T cells. **(b)** Schematic of the gene-program (GP) representation of T cell gene-expression profiles. Each cell is represented as a linear combination of gene programs, and each GP is defined by the upregulation or downregulation of a subset of genes. GPs are learned using EBMF, which decomposes the expression of gene *j* in cell *i* (*x*_*ij*_) as a sum over *K* gene programs, *x*_*ij*_ = *l*_*i*1_ *f*_*j*1_ + … + *l*_*iK*_ *f*_*jK*_, where *l*_*ik*_ is the activity level of program k in cell *i*, and *f*_*jk*_ is the effect of GP *k* on expression of gene *j*. **(c)** GP–gene network in which all 200 GPs are linked to their five most strongly regulated genes. Grey nodes are GPs, and tan nodes are genes. Selected GPs are highlighted in different colors, including the pan-T cell GP1, lineage-specific GPs such as GP58 in CD8 T cells and GP68 in Tregs, a resting-like GP171, helper T cell programs such as GP56, GP161, skin-associated GP6 and GP7, and the TCR-response GP35. **(d)** Heatmap showing the activity of the 200 GPs across T cells grouped by lineage and activation state. GPs are highlighted in colors corresponding to a subset of those used in (c).

To reveal the transcriptional structure underlying the atlas, we used two complementary approaches. We first applied EBMF^24^ and developed RQVI, detailed below, as an independent framework for assessing the robustness of the inferred GPs while reducing computation time from days to minutes. EBMF decomposes the gene expression matrix into gene programs and their cell-specific activity levels. In more detail, we decomposed the expression of gene *j* in cell *I* (*x*_*ij*_) as a sum over *K* gene programs, *x*_*ij*_ = *l*_*i*1_ *f*_*j*1_ + … + *l*_*iK*_ *f*_*jK*_, where *l*_*ik*_ is a non-negative number representing the activity level of program k in cell *i*, and *f*_*jk*_ is a number representing the effect of GP *k* on expression of gene *j* (**Fig. 1b**). Effects *f*_*jk*_ that are positive therefore capture upregulation of a gene, negative effects correspond to downregulation, and zeros represent no change. This “semi-NMF” decomposition is motivated by the idea that a gene program cannot be “negatively present” in a cell, but gene programs can involve both upregulation and downregulation (see also Liu et al.^25^). We applied the EBMF algorithm, implemented in “flashier”^31^, to identify up to K=200 GPs. To assess statistical replicability of the estimated GPs, we applied the same algorithm separately to each of the 80 sequencing batches (IGT) and computed cosine similarity between programs across replicates (see Methods). The majority of GPs were replicated in at least two batches, and the total number of replicated GPs plateaued at approximately 140, suggesting that increasing the number of GPs beyond 200 was unlikely to yield substantially more reproducible biological structure (**Extended Data Fig. 1a,b**).

Technical variation during data acquisition was minimized through shared standard operating procedures, single-experimentalist processing, centralized sequencing, and a unified data-processing pipeline. Nonetheless, some technical variation may remain, so we first assessed the remaining contribution of batch effects to GP activity. We leveraged the standard reference cells (spleen T cells from 6-8 week-old B6J mice) included in each experiment to quantify residual batch effects. Only a few GPs were associated with batch effects, and EBMF captured distinct forms of technical variation (**Extended Data Fig. 1c)**. For example, GP9 was enriched in specific batches (IGT13 and IGT14), whereas GP1 reflected a more continuous technical effect associated with sequencing depth (**Extended Data Fig. 1d,e; Extended Data Table 1**). Other sources of batch effect, including differences in tissue processing, may still be present, but these effects do not appear to be dominant, as no GP was specific to a single sample or tissue (**Extended Data Table 1**).

The network in **Fig. 1c** and heatmap in **Fig. 1d** provide a bird’s eye view of GP activity across lineages, and activation levels. Some GPs were broadly active across T cells and included large numbers of genes, consistent with core transcriptional programs shared by many T cells (**Extended Data Table 2**). GP1 was the clearest example: it was active in nearly all cells and captured the baseline, or mean, expression of many pan-T and housekeeping genes, including *Actb, Actg1, B2m*, TCR-associated genes, *Cd3d/e/g, Cd52* (**Fig. 1d,2a**). Another example was GP58, whose level of activity defined a quantitative axis separating more CD8-associated from more CD4-associated transcriptional states (**Fig. 1c,d)**. In contrast, most GPs were much more sparsely distributed, active in only a small subset of cells, often <1% of all cells (**Fig. 1d,2b**). These involved a smaller set of genes, typically fewer than 10-20 highly active genes per program (**Fig. 2c, Extended Data Fig. 1f**). Compared with alternative matrix factorization approaches, this sparsity is a key feature of EBMF and substantially improves GP interpretability^24,25^. Among these sparse GPs, some captured lineage-specific gene expression, such as GP68 in Tregs, whereas others represented transcriptional modules shared across lineages, including proliferation-associated programs (e.g. GP5, 21, 118), tissue-associated programs (e.g., GP6 in skin), or helper T cell-associated programs (e.g.,, *Il17*-enriched GP161 across CD4, gdT and Tz cells) (**Fig. 1c,d, and subsequent sections**). Altogether, each single cell expressed, on average, approximately 10 GPs **(Fig. 2d**), substantially simplifying the interpretation of gene expression patterns. The number of active GPs did not vary significantly across T cell lineages (**Fig. 2d**), but activated T cells expressed more GPs than resting cells (**Fig. 2e**), and the number of active GPs correlated with CD44 protein expression measured by CITE-seq (**Fig. 2f**).

**Figure 2.**
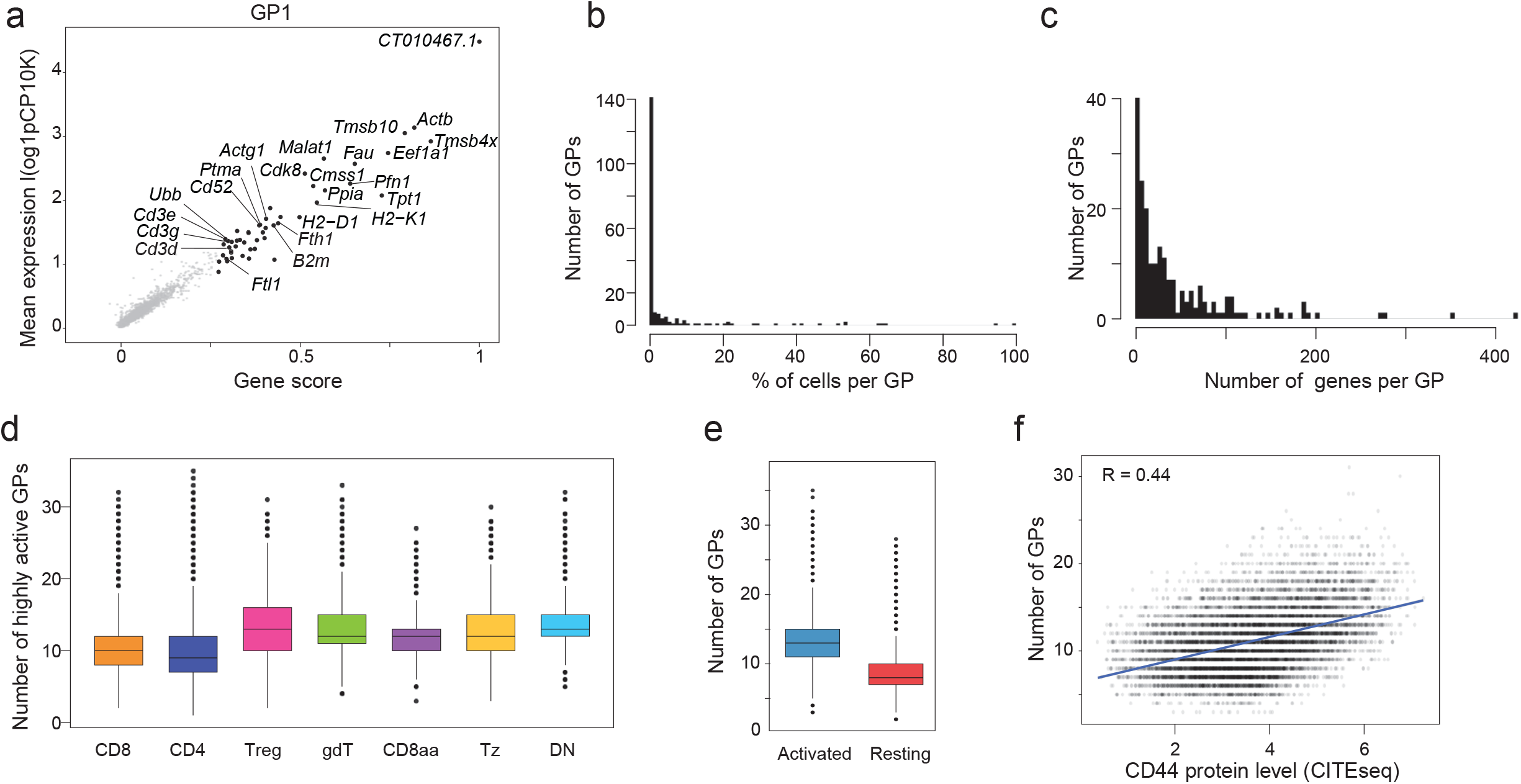
Individual T cells deploy sparse combinations of gene programs that expand with activation. **(a)** Gene scores for GP1, a baseline program capturing the pan-T cell transcriptional profile. Each point represents a gene. The x-axis shows the GP1 gene score, representing the magnitude and direction of gene regulation within the program and scaled to a maximum absolute value of 1. The y-axis shows mean expression across all T cells (log-normalized). **(b)** Histogram showing the fraction of cells in which each GP was highly active (loading greater than 0.1 in a given cell). **(c)** Histogram showing the number of highly active genes per GP (genes with an absolute score greater than 0.25 in a given GP). **(d)** Box plot showing the number of highly active GPs per cell across T cell lineages. **(e)** Box plot showing the number of highly active GPs per cell in activated and resting CD4+ and CD8+ T cells. **(f)** Scatterplot showing the number of highly active GPs per cell correlated with CD44 protein expression (CITE-seq; log-normalized counts). The blue line indicates the least-squares fit; Pearson’s r = 0.44.

Together, these results establish EBMF-derived gene programs as a sparse, interpretable, and reproducible layer of the immgenT atlas. In the following sections, we use this program-based representation to identify shared and context-specific transcriptional programs.

### Lineage and cluster-associated programs

The immgenT framework organizes T cells along several major axes of variation, including the separation of eight broad T cell lineages (“level 1”) and their clusters (“level 2”, e.g., CD4.A, CD8.B)^12^. We therefore asked how EBMF-derived GPs intersected with lineage-associated transcriptome structure. For each GP, we evaluated its sensitivity and specificity in predicting lineage identity, as summarized by the area under the curve (AUC) (**Fig. 3a, Extended Data Table 3**). An AUC close to 1 indicates a strong association between GP and lineage identity.

**Figure 3.**
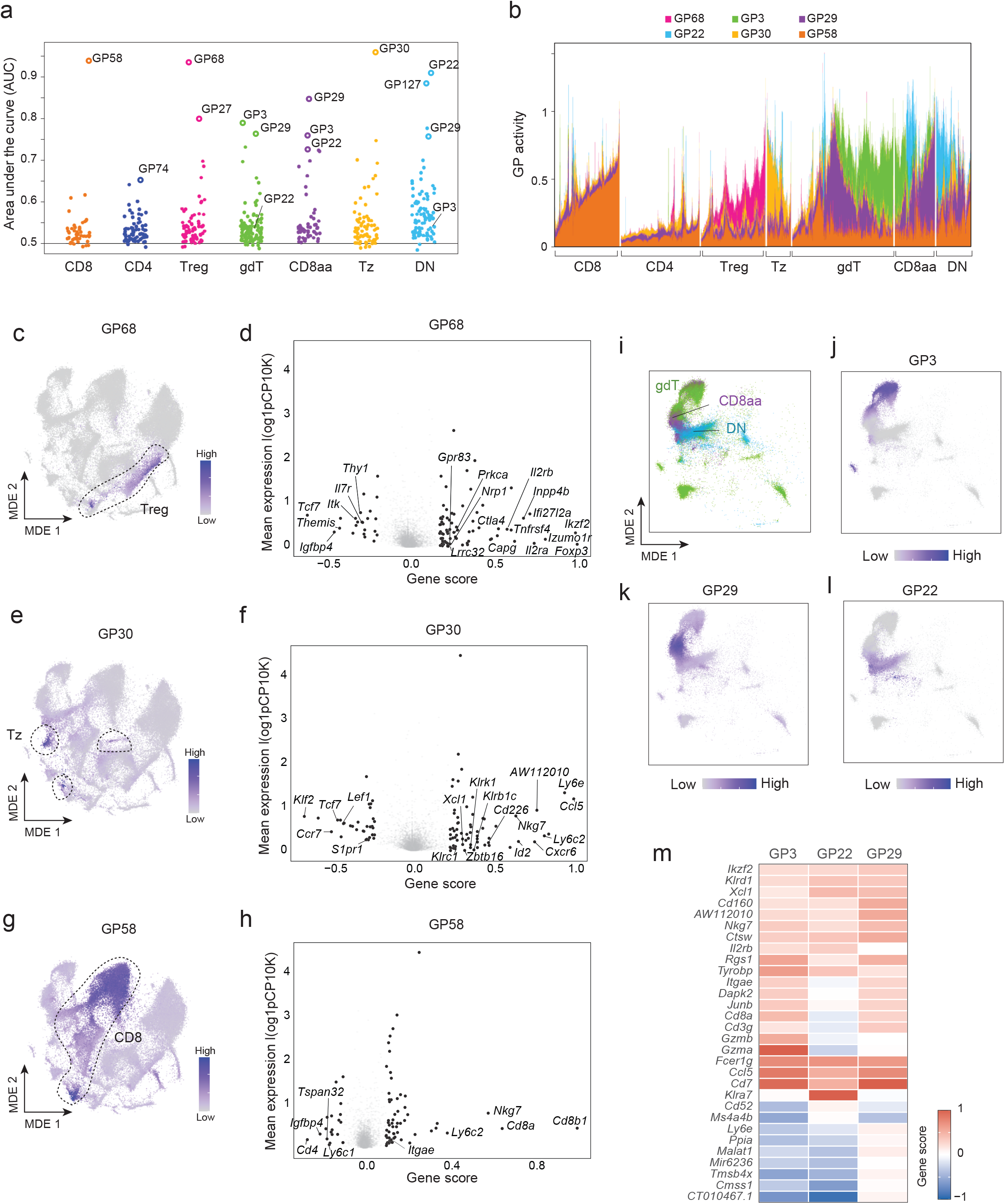
T cell lineage identity is encoded by both lineage-restricted and shared gene programs. **(a)** Swarm plot showing the AUC of each GP for predicting major T cell lineage. The top GPs for each lineage are highlighted. **(b)** Structure plot showing single-cell activity of the lineage-related GPs identified in (a), grouped by lineage. **(c)** All-T MDE colored by GP68 activity. **(d)** Gene scores for GP68. The x-axis shows the GP68 gene score, and the y-axis shows mean expression across all T cells (log-transformed). **(e)** All-T MDE colored by GP30 activity. **(f)** Gene scores for GP30, shown as in (d). **(g)** All-T MDE colored by GP58 activity. **(h)** Gene scores for GP58, shown as in (d). **(i-l)** All-T MDE subsetted to show only gdT cells, CD8aa T cells, and DN T cells. Cells are colored by **(i)** lineage (gdT in green, CD8aa in purple, and DN in blue), **(j)** GP3 activity, **(k)** GP29 activity, and **(l)** GP22 activity. **(m)** Heatmap showing the top 30 genes upregulated in GP3, GP22, and GP29.

The clearest examples of lineage-restricted programs were found for Tregs and Tz cells, marked by GP68 and GP30, respectively (**Fig. 3b**). GP68 was specifically expressed in Tregs and included canonical Treg-associated genes such as *Foxp3, Il2ra*, and *Ikzf2*, many of which are bound by *Foxp3*, as described in the immgenT Treg companion manuscript^28^ (**Fig. 3c,d**). GP30 was enriched in Tz cells and included *Zbtb16*, the central transcriptional regulator of several unconventional T cell lineages^32^ (**Fig. 3e,f**). This was notable because Tz clusters occupy distinct regions of the global MDE, in contrast to Tregs, which form a more continuous lineage structure. Thus, despite the heterogeneity of Tz populations, GP30 extracted a shared transcriptional program across these subsets (**Extended Data Fig. 2a**).

The most CD8-associated program, GP58, was led by *Cd8b1* and *Cd8a* and was most active in CD8αβ+ T cells (**Fig. 3g,h)**. However, GP58 also occurred in other lineages, if at a lower level. This likely reflects the contribution of cytotoxicity-associated genes such as *Nkg7, Ctsw*, and *Klrd1*, which are also expressed by CD8aa, gdT, DN, and subsets of CD4+ T cells. Interestingly, GP58 was more active in resting than activated CD8+ T cells (**Extended Data Fig. 2b**), capturing *Cd8a* and *Cd8b1* downregulation during activation, which we also observed by CITE-seq (**Extended Data Fig. 2c)**. Conventional CD4^+^ T cells did not show a comparably strong positive lineage-specific gene program. Instead, CD4 identity was best reflected by the absence of the CD8-associated GP58, for which *Cd4* was among the most negatively associated genes.

Three gene programs (GP3, GP22, and GP29) showed strong association with multiple lineages, gdT, CD8aa and DN. These lineages occupy partially overlapping regions in the MDE (**Fig. 3i**), and each GP captured a distinct region of overlap (**Fig. 3j-l**). The DN-enriched region was most distinct (GP22, AUC > 0.9), while GP3 and GP29 mapped onto regions shared by gdT and CD8aa cells. All three programs included *Cd7, Fcer1g, Ctsw*, and *Ccl5* (**Fig. 3m**), but differed in lineage-specific ways: *Cd8a* was reduced in GP22, and cytotoxic markers varied, with *Gzma*/*Gzmb* prominent in GP3 and killer-cell receptor genes in GP22.

Together, lineage-associated gene programs were rare and most clearly resolved for Tregs, Tz cells, and CD8αβ T cells. In contrast, gdT, CD8aa, and DN cells were organized by a set of related but distinct programs that mapped onto overlapping regions of the atlas.

We next examined GP organization across the 107 level 2 clusters defined within the eight T cell lineages^12^ (**Fig. 4a)**. GP activity was more predictive of level 2 cluster identity than of lineage identity, consistent with the greater transcriptional heterogeneity captured by these clusters (**Fig. 4b**). No level 2 cluster was defined by a single GP. Instead, each cluster was characterized by a distinct combination of programs, while individual GPs were shared across clusters and, in some cases, across lineages (**Fig. 4a, Extended Data Fig. 2d**).

**Figure 4.**
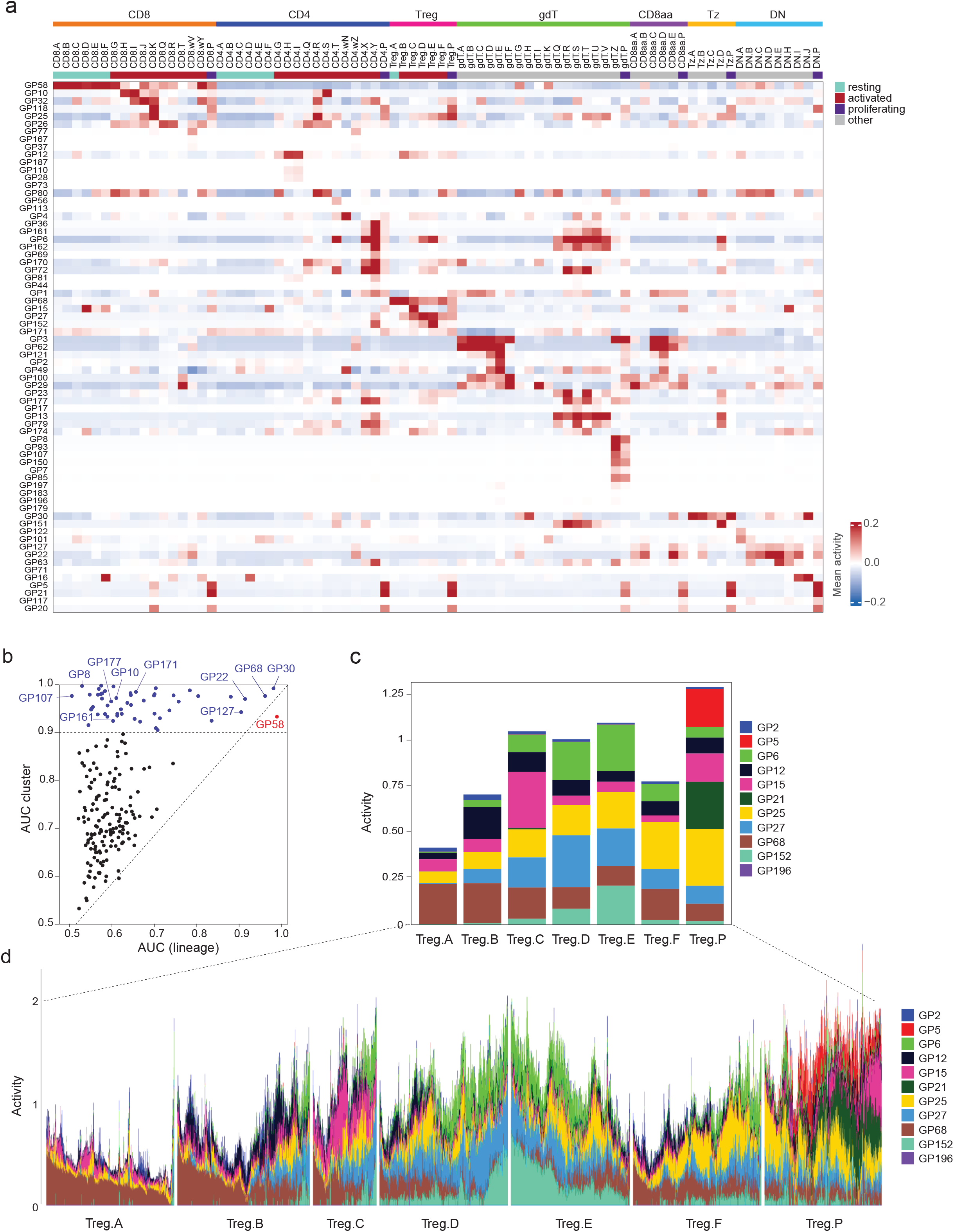
Shared transcriptional modules across T cell states. **(a)** Heatmap showing the row-centered mean GP activity across the 107 level 2 clusters for selected GPs with cluster AUC > 0.9, as identified in **a**. Mean GP activity was calculated in baseline samples. **(b)** Scatterplot comparing the predictive performance (AUC) of each GP for lineage identity (level 1; x-axis) and cluster identity (level 2; y-axis). GPs with AUC > 0.9 on either axis are highlighted and labeled (lineage-associated, red; cluster-associated, blue). **(c)** Stacked bar plots of the mean activity of the 11 GPs with AUC > 0.9 for at least one Treg cluster, illustrating that individual clusters are defined by distinct combinations of GPs rather than by a single cluster-specific program. **(d)** Structure plot showing single-cell activity of the 11 Treg-associated GPs shown in **c**, with cells grouped by Treg cluster. Variation in GP activity within clusters illustrates transcriptional heterogeneity not captured by discrete cluster assignments.

Treg clusters provided an illustrative example. They were distinguished by different combinations of a relatively small set of 11 GPs, some of which showed strong cluster associations, such as GP21 in proliferating Tregs and GP15 in Treg.C. Other clusters shared much of their GP composition but differed quantitatively in the activity of individual programs (**Fig. 4c**). For example, Treg.D and Treg.E shared most of their active GPs, but GP27 was more prominent in Treg.D, whereas GP152 was more active in Treg.E. Because GP activity is defined at single-cell resolution, the representation also captured heterogeneity within clusters (**Fig. 4d)**. Within Treg.E, for example, GP25 and GP152 varied substantially among cells and were inversely associated. Thus, the GP representation recapitulated cluster structure while offering complementary advantages: it revealed transcriptional commonalities across clusters and captured heterogeneity within clusters.

### Gene programs in activated CD4+ and CD8+ T cells

Conventional CD4^+^ and CD8αβ T cells exhibit broad functional diversification upon activation, encompassing effector, cytokine-response, exhaustion, tissue-location, and proliferation-associated programs^33,34^. We therefore investigated how gene programs organize these activated states. Comparing GP activity in activated CD4+ and CD8+ T cells with their resting counterparts as defined by CD62L and CD44 expression (**Extended Data Fig. 3a**) revealed widespread activation-induced remodeling of GP activity (**Fig. 5a**): 40 and 25 programs showed a mean fold change greater than 3 relative to resting CD4+ and CD8+ T cells, respectively (**Extended Data Table 4**). Only six programs were downregulated, most of which were shared between lineages. These included GP171, a resting/recirculating program marked by *Ccr7, Lef1, Tcf7*, and *Klf2*, together with additional resting-associated programs reflecting heterogeneity within the resting compartment (**Fig. 5b, Extended Data Fig. 3c, Extended Data Table 2**).

**Figure 5.**
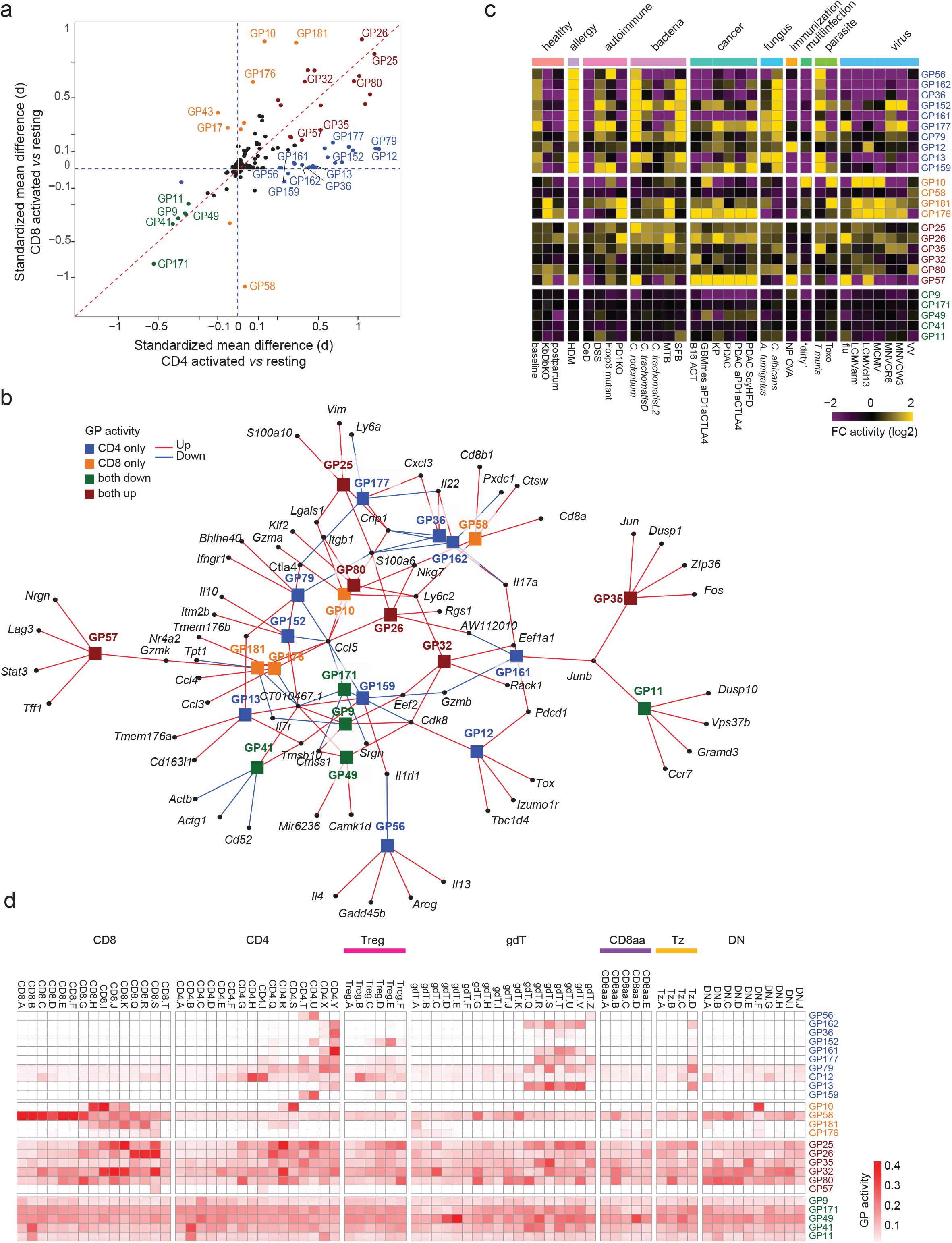
Activated T cell states arise from shared and lineage-biased combinations of gene programs. **(a)** Standardized mean difference in GP activity between activated and resting cells in CD4+ T cells (x-axis) and CD8+ T cells (y-axis). Points are colored according to whether they are highly upregulated in both CD4+ and CD8+ activation (brown), downregulated in both (green), or preferentially upregulated in CD4 (blue) or CD8 (orange). **(b)** GP-gene network in which each GP is linked to its five most strongly regulated genes, based on GP gene scores. Edges are colored by the sign of the gene score (red for upregulated, blue for downregulated). GP nodes are colored as in (a). **(c)** Heatmap of log2 fold change in mean GP activity across experimental conditions for activated CD4+ and CD8+ T cells, computed relative to each GP’s mean loading across all CD4+/CD8+ cells. Columns grouped and annotated by immune challenge category. GPs are ordered and colored as in (a). **(d)** Heatmap of mean GP activity per level 2 cluster, with the top bar denoting parent lineage. GP ordered and colored as in (a).

Among activation-induced programs, 16 were shared between CD4+ and CD8+ T cells. GP26 and GP80 were the most active, present in 86% of activated T cells, and associated with programs of tissue localization and recirculation. GP26, characterized by reduced *Klf2* expression, a critical regulator of tissue exit^35^, and increased *Rgs1* expression^36,37^, was enriched in non-lymphoid tissues and broadly active in activated T cells from organs such as the prostate and submandibular gland (**Fig. 5b; Extended Data Fig. 3b**). In contrast, GP80 captured a *Klf2*-associated recirculation program marked by integrin expression. Another shared program, GP35, represented an immediate TCR-response state characterized by *Nr4a1, Fos, Jun*, and *Dusp1* expression^38,39^ (**Fig. 5b, Extended Data Fig. 3c**).

Nine programs were preferentially induced in activated CD8+ T cells, including GP10, marked by cytotoxic effector genes such as *Gzma* and *Nkg7*, and GP181, led by *Gzmk* (**Fig. 5b**), a protease that does not trigger cell death but instead activates the complement pathway^40^. In contrast, 24 programs were preferentially induced in activated CD4+ T cells. Many corresponded to helper T cell functions, although these were often distributed across multiple gene programs. For example, the Th2-like response was split between GP56, marked by *Il4, Il13*, and *Areg*, and GP159, which captured *Il1rl1* expression^41^. Several programs were associated with IL-17 expression, including GP13, GP36, and GP161^42^. Among these, GP36 showed co-expression of *Il22*. Another program, GP162, captured *Il22* expression independently of *Il17*, suggesting partially distinct regulation of IL-22 production^43^. GP152 exhibited a more regulatory profile, characterized by expression of *Ctla4* and *Il10*.

Additional CD4-biased programs captured potentially novel activation modalities. GP79 was characterized by *Odc1, Bhlhe40, Ifngr1, Il7r, Rgs1, Il18ra, Rora*, and *Tmem176a*/*b*, and was broadly detected across diverse inflammatory settings (**Extended Data Fig. 3d**). This program was shared among activated CD4 populations spanning multiple Th phenotypes (**Extended Data Fig. 3e**). GP177, marked by *Cxcl3*, a neutrophil chemoattractant^44^, was specific to CD4 cells and particularly enriched in cells from CD4.X/Y, the most polarized Th17-like populations (**Fig. 5b, Extended Data Fig. 3e)**.

Activated CD4+ and CD8+ T cells also shared chronic-stimulation programs. GP57 contained genes that have been associated such as *Tox* and *Lag3* and was active in both CD4+ and CD8+ tumor-infiltrating T cells^45,46^ (**Fig. 5c; Extended Data Fig. 3f; Extended Data Table 2**). However, chronic activation also exhibited lineage-biased structure. GP12 was preferentially active in CD4+ T cells and was characterized by expression of *Izumo1r* (FR4)^47^ and *Pdcd1*, whereas GP176 was preferentially active in CD8+ T cells and captured increased expression of *Nr4a1, Nr4a2, Lag3, Tigit, Tox*, and *Pdcd1*.

Activation GPs were increased during challenges (**Fig. 5c**) and were often shared across multiple conditions. For example, Th2 GP56 was active during *T. muris* infection and was also observed in *Foxp3*-deficient mice, consistent with the well-established Th2 skewing seen in such mice^48^. Likewise, both influenza infection and cancer were associated with activation of GP177, indicating that T cells can deploy common gene programs across diverse contexts. This recurrence across unrelated perturbations supports the interpretation of GPs as shared transcriptional modules, rather than signatures of individual experiments or isolated disease settings. Correspondingly, individual conditions were typically characterized by multiple activation programs rather than a single Th-associated GP. For example, *T. muris* infection led to activation of both the Th2-associated GP56 and Th17-associated GP36 programs.

At single-cell resolution, GPs were enriched in specific immgenT clusters but were rarely cluster-exclusive, supporting a quantitative view in which clusters represent regions of high program activity rather than discrete transcriptional identities (**Extended Data Fig. 3e**). Moreover, programs active in conventional CD4+ or CD8+ T cells were active across other lineages **(Fig. 5d**). For example, the cytotoxic GP10 was active across CD8.H-K, CD4.S, and DN.F clusters. A Th2-associated program, GP56, was present in CD4.T/U cells but also in gdT cell cluster gdT.Z, whereas the Th17-associated GP36 was enriched in CD4.X/Y and also in gdT.T/U. Similarly, a Tfh-associated GP12 was active in both CD4.H and Treg.C, where Tfh and Tfr map, respectively.

Thus, activated T cell states are organized by shared transcriptional modules that are deployed in lineage- and context-specific combinations. This program-based view was particularly informative for states of CD4+ T cells. As discussed in the companion immgenT-CD4 manuscript^26^, Th1/2/17 nomenclature did not correspond to separate clusters, and most activated cells occupied mixed clusters with low-level combinations of Th-associated signatures. Our results further show that there is no simple one-GP-per-Th-subset model. Instead, each Th-associated state was represented by several distinct GPs, whose combinations varied across clusters and lineages.

### Tissue-associated gene programs are often deployed in specific T cell subsets

T cells from different tissues show marked transcriptional differences, which may reflect either differences in the T cell populations present in each tissue or tissue-imposed programs acting across multiple populations^49,50^. The companion immgenT-Cosmology manuscript showed that tissues differ substantially in their T-cell cluster composition. Here, we revisited tissue-associated variation from a GP perspective. Thirty-two GPs showed differential activity across tissues, with each tissue characterized by a distinct combination of programs (**Fig. 6a**). Seven were highly predictive of individual tissues (AUC > 0.9; **Extended Data Table 5**). These tissue-associated programs were shared across multiple T cell lineages (**Fig. 6b**). For example, GP37 was broadly active across mammary gland T cells (**Fig. 6c**). GP26, described above, represented a shared non-lymphoid tissue program characterized by *Rgs1* expression and reduced *Klf2*, consistent with tissue residency^35,51,52^.

**Figure 6.**
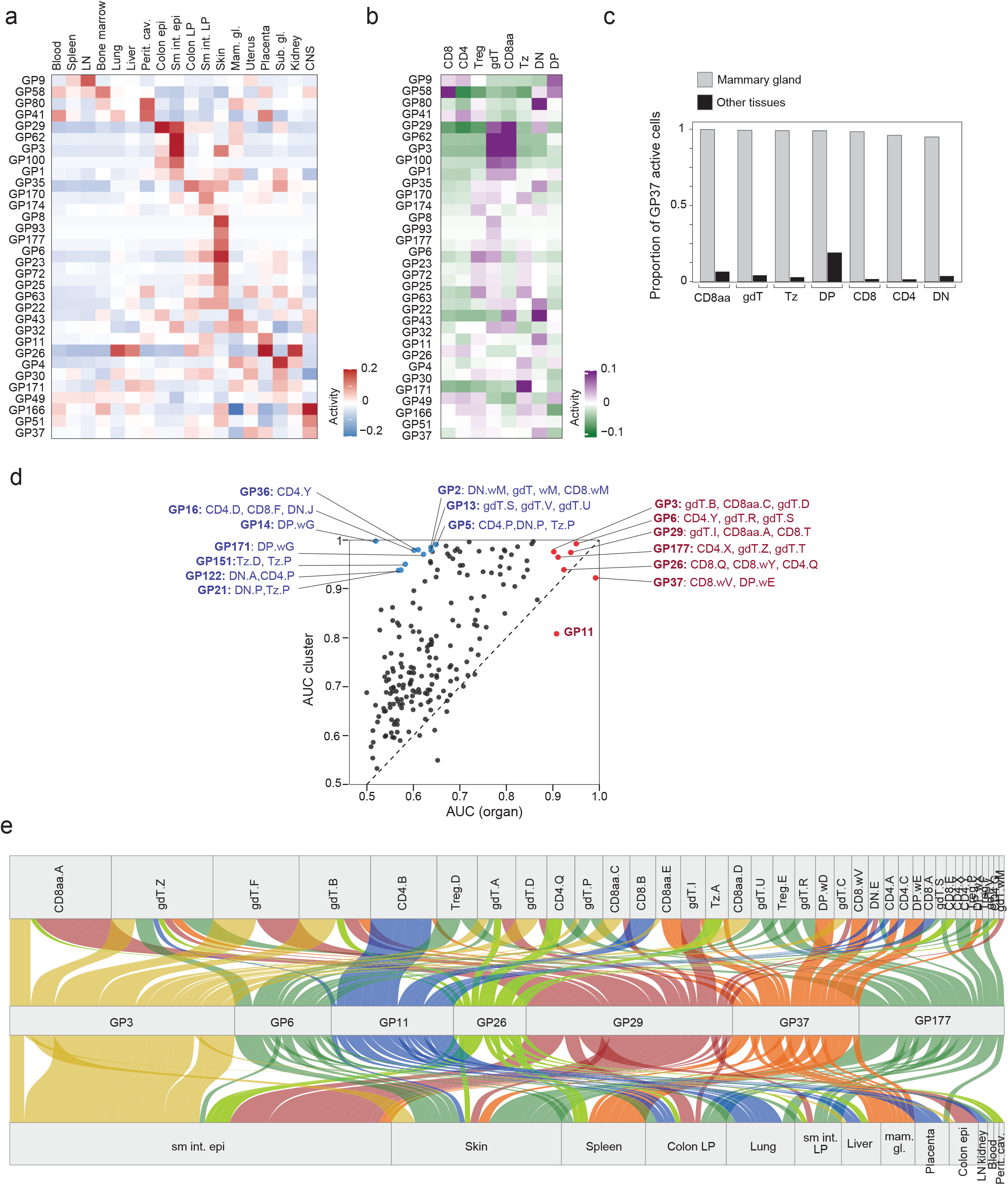
Tissue-associated gene programs reflect both tissue adaptation and T cell subset composition. **(a)** Heatmap showing the row-centered mean GP activity across 18 tissues for the 32 GPs with a tissue-specific mean loading > 0.1 or an AUC > 0.9. Mean GP activity was calculated in baseline samples. **(b)** Heatmap showing the row-centered mean activity of the same 32 GPs across T cell lineages. Mean GP activity was calculated in baseline samples. **(c)** Proportion of GP37-active cells in each lineage in the mammary gland compared with all other organs. **(d)** Scatterplot showing the predictive performance (AUC) of each GP for tissue of origin (x-axis) versus cluster identity (y-axis). Seven tissue-specific GPs with an AUC > 0.9 for tissue of origin are shown in red. Cluster-specific GPs with low organ-AUC are shown in blue. **(e)** Alluvial plot showing how tissue-specific GP activity is distributed across clusters. For each organ, GP-active cells are traced to their cluster annotation. Flows are colored by GP, and width is proportional to the number of cells.

These 32 tissue-associated GPs were, perhaps surprisingly, more strongly associated with level 2 cluster identity than with tissue identity, with the GPs lying above the identity line in **Fig. 6d**. The seven GPs with the strongest tissue associations were also highly predictive of specific level 2 clusters (AUC > 0.9), indicating that tissue-associated programs were preferentially deployed within particular T cell states. The alluvial plot in **Fig. 6e** illustrates several key features of how tissue and cluster relationships coexist. First, tissue-associated GPs stand out clearly, but none were absolutely tissue-restricted. For example, GP3 was most prominent in the small-intestinal epithelium but could also be detected in a related gdT cell cluster in the skin. Second, and in keeping with Fig. 6d, these tissue-preferential GPs were predominantly expressed in defined clusters within each lineage. GP37, a mammary gland-associated program, was expressed across multiple lineages, but within each lineage it was preferentially used by cells of one cluster. This cluster-level organization could take two forms. In some cases, the same GP mapped to analogous clusters across multiple tissues, consistent with shared cluster identities contributing to shared gene expression. For example, GP11 was expressed primarily in CD4.A/B and CD8.A/B cells across lung, liver, placenta, kidney, colon lamina propria, and skin. In other cases, the same GP mapped to different clusters depending on tissue and lineage. GP177, for instance, was active in Treg.F in the placenta, in Treg.E in the colon, and in CD4.X/Y cells in the small intestine.

Thus, the GP is consistent with the orthogonal analysis in the immgenT cosmology manuscript^12^, which showed shared clusters across tissues together with tissue-specific differences in cluster composition. It further shows that tissue-associated GPs do exist, and are preferentially deployed in specific clusters.

### Mapping surface protein expression onto gene programs

The immgenT data included 128 surface markers evaluated by CITEseq in the same cells, allowing us to relate surface-protein expression to GP activity. We asked whether combinations of these markers, many of which are commonly used by for flow cytometry, could serve as proxies for specific transcriptional programs. To do so, we projected CITE-seq protein expression onto the RNA-defined GP space using semi-non-negative EBMF, fixing the cell-loading matrix learned from gene expression to estimate a protein signature for each GP (see Methods and **Fig. 7a**). The protein-GP association heatmap revealed broad combinatorial. Many of the markers were individually asociated with many GPs (e.g. CD44 alone associated with 89 programs; **Extended Data Fig. 4**). This many-to-many relationship was compounded by context-dependence: the same marker could report different programs across lineages or activation context, as illustrated by KLRG1 and CD69. KLRG1, expressed in subsets of CD4, CD8, and Treg cells, was associated with GP10 in CD4^+^ and CD8^+^ T cells (**Fig. 7b**), consistent with cytotoxic or CX3CR1^+^ Th1 effector states^53^, but with GP27 in Tregs, a distinct program associated with effector Treg functions and non-cytotoxic genes (*Areg, Tnfrsf4*)^28^ (**Fig. 7c, Extended Data Table 2**). Compared to KLRG1, CD69 associations varied across tissues and activation states rather than lineages **(Fig 7d, Extended Data Fig. 5a-b)**. In non-lymphoid tissues, CD69 correlated with a spectrum of NLT-associated programs: GP26 and GP63, broadly active across T cell lineages in NLT; GP3, GP29, and GP62 in gut-resident CD8aa intraepithelial T cells; and GP6 in skin, marked by *Vim, Lgals1*, and *S100a6*. These associations are consistent with the role of CD69 in sequestering S1PR1 and preventing tissue egress^54^. By contrast, transient CD69 induction during early T cell activation^55^ tracked with the TCR-response program GP35 and was anti-correlated with the resting program GP171.

**Figure 7.**
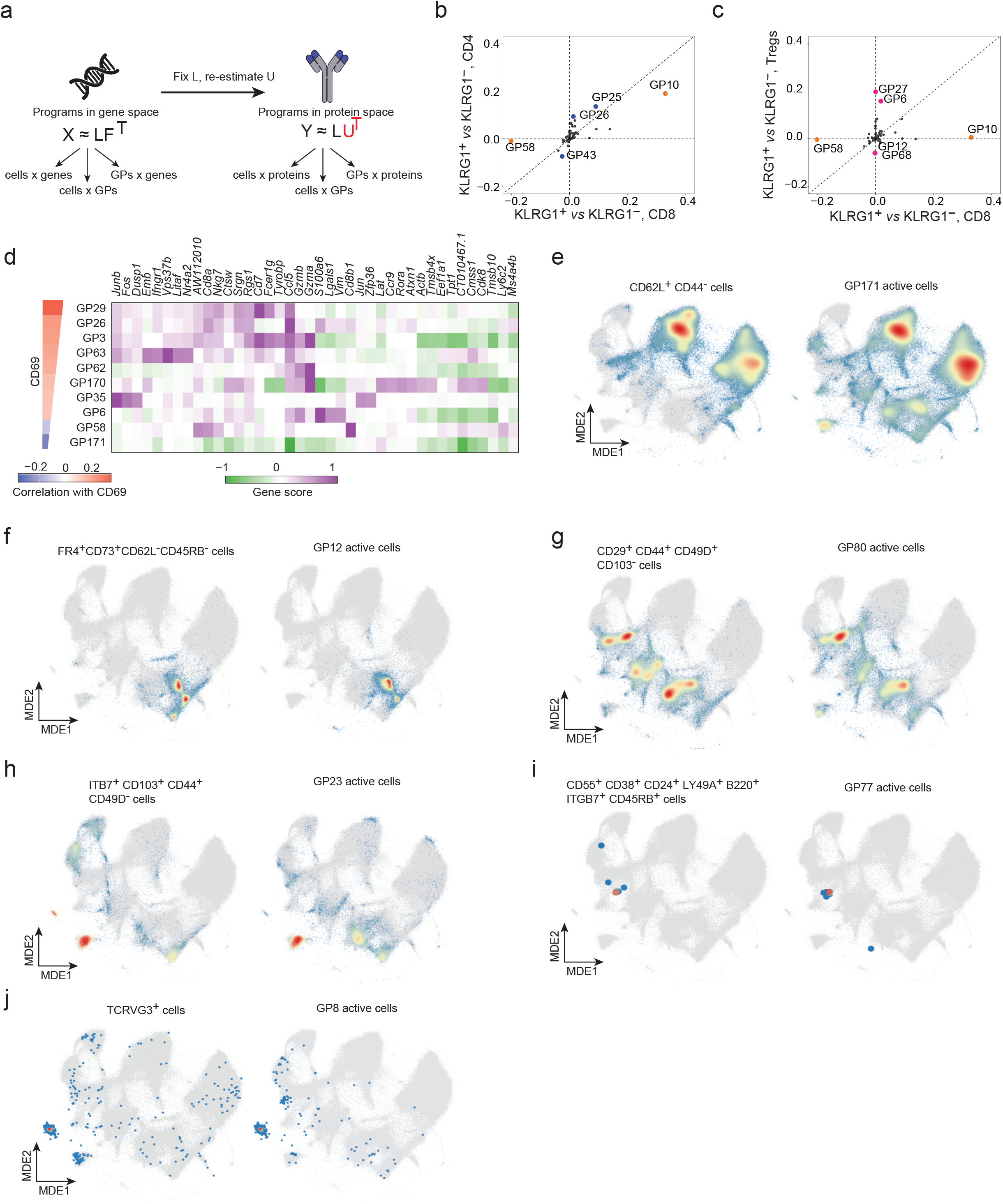
Surface-marker associations are context-dependent, but combinatorial gating resolves specific gene programs. **(a)** Schematic illustrating how protein signatures were defined for each GP. Holding the cell-loading matrix *L* fixed from the GP model, a protein factor matrix was estimated by EBMF from the paired CITE-seq protein measurements, *Y* ≈ *LU*^T^ (cells × proteins matrix), yielding a protein signature *U* (proteins × GPs matrix) for each GP. **(b-c)** GPs associated with KLRG1 protein expression in CD4, Tregs, and CD8. Effect-size versus effect-size plots, where each GP’s effect size is the difference in its mean activity between KLRG1^+^ and KLRG1^-^ cells, comparing CD8^+^ T cells (x-axis) with CD4^+^ T cells (b) or Treg cells (c). **(d)** Heatmap of the top five most strongly up- and down-regulated gene scores for ten GPs correlated with CD69 expression, with the left strip indicating the Spearman correlation between GP activity and CD69 expression. **(e-j)** Examples of gating strategies used to identify GP-active cells. All-T MDE plots highlight cells selected by the proposed gating strategy (left) and the corresponding GP-active cells (right). Color indicates cell density.

Marker combinations yielded greater specificity, correctly resolving lineage-associated GPs as expected (GP29, GP58, GP22, GP68; **Extended Data Fig. 5c-f**). More notably, the same GP was in several cases shared across lineages and recovered by the same protein combination, demonstrating that combinatorial gating can report conserved transcriptional states. GP171, active in resting CD4, CD8, and Treg cells, corresponded to the canonical CD62L^+^ CD44lo phenotype (**Fig. 7e**). FR4 and CD73 together marked the “anergic” program GP12 across CD4 and Tregs^56^ (**Fig. 7f**). Shared integrin combinations delineated common programs: CD29^+^ CD49D^+^ CD103^−^ ITGB7^−^ cells captured GP80, a *Klf2*-dependent tissue-egress program active across Tregs^28^, CD4, CD8, and DN cells (**Fig. 7g**), whereas CD103^+^ ITB7^+^ CD29^−^ CD49D^−^ cells marked the tissue-residency program GP23 across lineages (**Fig. 7h**). We also recovered rarer programs, including GP77 in B220^+^ CD8^+^ mammary gland T^57^ cells and GP8 in DETCs^58^, identified by TCRVγ3 and TCRγδ expression (**Fig. 7i,j**).

In conclusion, no single marker uniquely identifies a GP, and the same marker can reflect distinct transcriptional states across lineages and contexts. Marker combinations, however, can recover specific transcriptional programs, including programs conserved across lineages, providing a basis for prospective isolation and functional study.

### RQVI, a deep learning framework, validates EBMF-derived gene programs

Recent advances in deep learning have enabled single-cell models to capture complex, non-linear patterns of gene expression. We therefore asked whether the 200 EBMF-derived gene programs were robust to a distinct modeling framework, and whether a non-linear approach would recover similar gene programs. To address this, we developed Residual Quantized Variational Inference (RQVI), a deep learning framework designed to extract interpretable gene programs from the immgenT latent space (**Fig. 8a**). RQVI starts from an scVI encoder^59^ that projects cells into a latent space, capturing T cell heterogeneity across the atlas. To interpret this latent space in terms of gene expression programs, RQVI extends the model with a discrete dictionary of gene programs^60^. In this framework, and in line with the biological schema of **Fig. 1b**, each cell is represented as a combination of programs, analogous to the loading matrix, and each program is represented by a weighted set of genes, analogous to a factor matrix. The model is optimized so that the reconstructed cell representation remains as close as possible to the original latent space, thereby preserving the structure learned by scVI while decomposing it into interpretable transcriptional components. RQVI learns feature maps that consist of discrete codes and recursively quantizes the feature map with the goal of learning finer representations in each iteration^61^ . RQVI also offered practical advantages. It trained in minutes on the full ∼700,000-cell immgenT dataset, compared with the week-long runtime required by Flashier on the same dataset (**Extended Data Fig. 6a**). Similar to EBMF, RQVI also produced sparse representations in both gene and cell space, facilitating interpretation of both program gene content and cell-level activity (**Extended Data Fig. 6b**).

**Figure 8.**
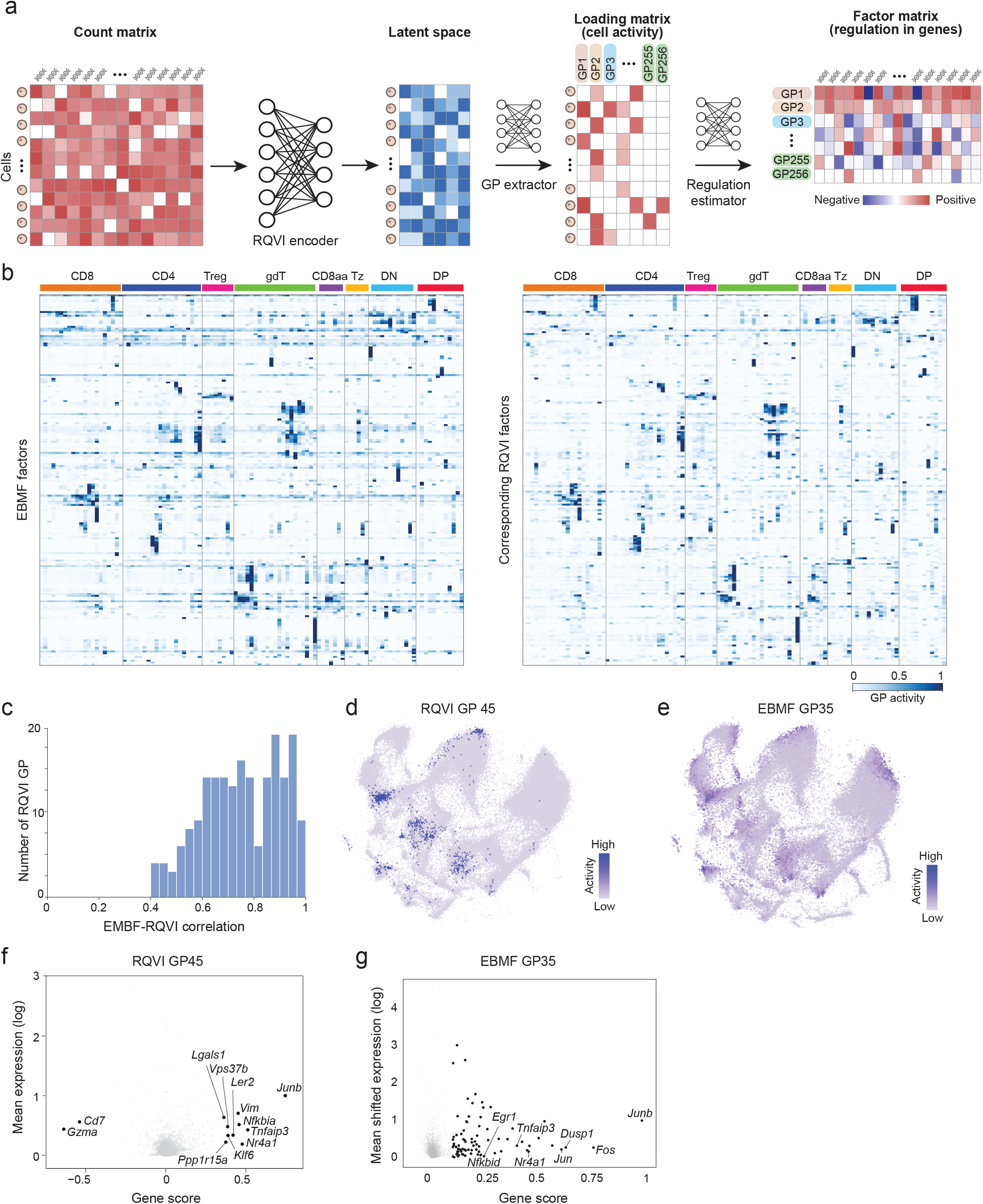
A nonlinear deep-learning framework independently recovers the immgenT gene-program landscape. **(a)** Schematic of the Residual Quantized Variational Inference (RQVI) framework. RQVI uses an scVI-style encoder to map cells into a continuous latent space, then reconstructs this representation through a discrete dictionary of reusable components. In effect, the model is asked to explain each cell’s position in the atlas using a limited set of program-like elements, rather than leaving the latent space as an uninterpreted coordinate system. The residual structure allows broad and more specific sources of variation to be represented within the same framework, which is biologically appealing given the layered nature of T cell identity, activation, tissue adaptation, and context-specific responses. Each component can then be scored across cells with the GP extractor and decoded into gene weights with the regulation estimator, yielding outputs directly comparable to EBMF gene programs. **(b)** Cluster-level activity profiles of corresponding EBMF and RQVI GPs. EBMF cell loadings (left) and matched RQVI program loadings (right) were averaged within the 107 level-2 clusters. **(c)** Histogram showing, for each EBMF GP, the best-matched Pearson correlation with any RQVI GP across 10 independent RQVI runs. **(d,e)** All-T MDE plots showing cell-level activity of a representative lower-correlation matched pair: (d) RQVI GP45 and (e) EBMF GP35. Color indicates GP activity from low to high. **(f,g)** Gene scores for RQVI GP45 (f) and its EBMF equivalent GP35 (g). The x axis shows the GP gene score, and the y axis shows mean expression across all T cells (log transformed).

Comparing GP activity between EBMF and RQVI across clusters revealed high agreement between the two approaches (**Fig. 8b**). To quantify this convergence, we calculated Pearson correlations between EBMF and RQVI program activity profiles across 10 independent RQVI runs and assigned each EBMF program a distinct RQVI match to maximize the overall correlation across all matched pairs (see Methods) (**Fig. 8c**). All 200 EBMF-derived gene programs correlated with at least one RQVI program, with a median best-match correlation of 0.75 and 95% exceeding r = 0.5. Program recovery also increased with the number of RQVI seeds. A single RQVI run recovered 79.5% of EBMF programs at |r| ≥ 0.5, whereas aggregating across seeds using a greedy set-cover strategy progressively recovered sparser and lower-signal programs that could be missed by any individual run. Thus, individual RQVI runs captured most of the gene-program landscape, while multi-seed aggregation, made practical by RQVI’s speed, improved recovery of subtle programs.

Further confirming the agreement between methods, the four lineage-associated EBMF programs (GP68 in Tregs, GP58 in CD8αβ T cells, GP30 in Tz cells, and GP22 in DN cells) had closely matched RQVI counterparts, with Pearson correlations of 0.94, 0.84, 0.91, and 0.90, respectively. These matched pairs showed concordant cell-level loading patterns and consistent leading genes (**Extended Data Fig. 6c-j**). Furthermore, even the lowest-correlated program pairs were biologically concordant. For example, RQVI GP45 and EBMF GP35 (r = 0.44) were active in the same cells and shared leading genes characteristic of a TCR-activation signature, including *Nr4a1, Fos, Jun*, and *Dusp1* (**Fig. 8d-g**). Thus, the differences in correlation magnitude didn’t reflect fundamentally distinct biological signals.

In conclusion, the 200 EBMF-derived gene programs were robustly recovered using an independent non-linear deep learning framework. Given the breadth and saturation of the immgenT atlas across mouse T cell lineages, organs, and immune challenges, this collection represents an extensive reference of gene programs underlying mouse T cell heterogeneity.

## Discussion

The immgenT GP analysis adds an interpretable layer beneath the cellular-state framework defined in the companion immgenT manuscripts^12,26–28^. T cell heterogeneity can be understood not only as a finite set of recurrent cell states, but also as a collection of gene programs deployed in different combinations across lineages, tissues, and immune challenges^5^. Our framework enables this perspective to be explored systematically across the breadth of the immgenT dataset. The 200 GPs are interpretable, and reference-free, independent of known labels or preselected signatures. Their strong convergence with programs inferred using the non-linear RQVI framework further supports their robustness. Some GPs correspond to major lineages, as expected, but lineage-specific GPs were not the rule. Tissue effects showed a different organization: tissue-associated GPs were readily detected but were most often deployed through specific clusters.

GP analysis provides a cluster-free framework for interpreting single-cell RNA-seq data^5,21,62,63^. Because each program is defined by the coordinated expression of multiple genes, it helps overcome single-cell sparsity and enables program activity to be studied at single-cell resolution. This reveals heterogeneity within clusters and captures shared modules and quantitative variation, which was particularly insightful in the context of helper-associated programs. In the companion immgenT CD4 manuscript^26^, T helper programs cut across clusters: rare highly polarized Th1, Th2, Th17, and Tfh states coexisted with dominant activation states expressing mixed and graded combinations of these programs. Here, we further found that helper-associated signals could themselves be partitioned into multiple related programs, enabling stacked or combinatorial expression within individual cells. For example, Th2-like features were divided between GP56 and GP159, whereas IL-17- and IL-22-associated expressions were distributed across multiple programs. These programs also extended beyond conventional CD4+ T cells and were detected across multiple T cell lineages, emphasizing their shared nature. Similar relationships were observed outside helper-associated states: GP62, for example, was highly active in CD8aa.C and CD8aa.D but largely absent from CD8aa.A and CD8aa.B, highlighting a shared *Gzma*/*Gzmb*-associated cytotoxic program among distinct clusters of gut-resident CD8aa T cells. Together, these observations may help explain why the same labels can carry different meanings across studies. Terms such as “exhausted,” “resident,” “cytotoxic,” and “Th1-like” often refer to combinations of programs and clusters rather than to single states or individual GPs.

This work provides a resource for both experimental and computational studies. First, GP activity across more than 700 immgenT samples can be explored through a dedicated website (see Methods). Second, the matrix-factorization framework underlying the 200 immgenT GPs provides a natural basis for projecting external datasets onto this reference, for example by fixing the learned gene-program matrix and estimating program loadings in new data. Robust application of this strategy will, however, require careful treatment of cross-dataset technical variation and recognition of cases in which additional programs are needed to capture biological states absent from the reference^23^. Third, the GP signatures provide a resource complementary to the Molecular Signatures Database (MSigDB^64^) for gene-set enrichment analyses such as GSEA^65^. Fourth, CITE-seq can be used to determine which GPs are associated with surface phenotypes in different cell types and conditions, as illustrated by KLRG1, which was linked to distinct programs in CD4+, CD8+, and Treg cells.

GPs denote recurrent patterns of covarying gene expression across cells and organize biological variation, but they do not by themselves define causal regulatory networks^14^. In some cases, a GP may closely reflect a coherent regulatory module, as exemplified by the Treg-associated GP68, whose leading genes overlap strongly with the canonical Foxp3-dependent Treg program^66^. In many, and perhaps most, cases, however, a GP may instead reflect partially independent regulatory circuits that converge because of shared upstream signals or other correlated biological processes. These limitations define the next step from descriptive gene programs to predictive models of gene regulation. Emerging AI-based and statistical approaches are increasingly being developed to infer regulatory logic from DNA sequence, chromatin accessibility, single-cell expression, and perturbation data. The immgenT GP atlas provides a useful resource for such approaches.

In conclusion, these analyses show that T cell heterogeneity reflects both discrete cellular states and recurrent transcriptional programs across tissues and conditions. ImmGenT-GP provides a systematic reference for resolving these complementary dimensions of T cell variation and for testing their conservation in human datasets.

## Acknowledgments

We thank the many colleagues who were consulted at various stages of this project. This work was funded by a grant from the NIH-NIAID to the ImmGen consortium (AI072073).

## Author Contributions

ZZ, SSP, TW, PC performed the computational processing and analyses; DZ, MS, SM, MB, CB and immgenT principal investigators designed the overall study; DZ, CB and immgenT principal investigators oversaw the experiments; ZZ, DZ, TW, MS, CB, MB wrote the manuscript with input from other authors.

## Collaborators

Participants in the immgenT Project include: Aaron Liu^1^, Alexander Chervonsky^2^, Alexandra Cassano^2^, Alia Welsh^3^, Amir Ferry^4^, Ananda Goldrath^4^, Andrea Lebron-Figueroa^5^, Ankit Malik^2^, Anna-Maria Globig^6^, Antoine Freuchet^2^, Bana Jabri^2^, Charlotte Imianowski^7^, Christophe Benoist^5^, Claire Thefaine^8^, Dan Kaplan^7^, Dania Mallah^5^, Dario Vignali^7^, David Sinclair^5^, David Zemmour^2^, Derek Bangs^9^, Domenic Abbondanza^2^, Enxhi Ferraj^10^, Eric Weiss^7^, Erin Lucas^8^, Evelyn Chang^10^, Gavyn Chern Wei Bee^11^, Giovanni Galletti^4^, Ian Magill^5^, Iliyan D. Iliev^12^, Joonsoo Kang^10^, Jordan Voisine^2^, Josh Choi^5^, Julia Merkenschlager^13^, Jun R. Huh^5^, Katharine Block^8^, Ken Cadwell^11^, Kennidy K. Takehara^4^, Kevin Osum^8^, Laurent Brossay^14^, Laurent Gapin^15^, Liang Yang^5^, Lizzie Garcia-Rivera^1^, Marc K. Jenkins^8^, Maria Brbic^16^, Maria-Luisa Alegre^2^, Marion Pepper^9^, Mariya London^17^, Matthew Stephens^2^, Maurizio Fiusco^16^, Melanie Vacchio^3^, Michael Starnbach^5^, Michel Nussenzweig^13^, Mitch Kronenberg^18^, Myriam Croze^19^, Nalat Siwapornchai^5^, Nathan Morris^12^, Nicole E. Scharping^4^, Nika Abdollahi^19^, Nitya Mehrotra^2^, Odhran Casey^5^, Olga Barreiro del Rio^5^, Paul Thomas^20^, Peter Carbonetto^2^, Remy Bosselut^3^, Rocky Lai^10^, Sam Behar^10^, Sam Borys^14^, Sara E. Hamilton^8^, Sara Mostafavi^9^, Sara Quon^4^, Serge Candéias^21^, Shanelle Reilly^14^, Shanshan Zhang^5^, Siba Smarak Panigrahi^16^, Sofia Kossida^19^, Stefan Muljo^3^, Stefan Schattgen^20^, Stefani Spranger^22^, Steve Jameson^8^, Susan M. Kaech^1^, Takato Kusakabe^12^, Taylor Heim^22^, Tianze Wang^9^, Tomoyo Shinkawa^10^, Ulrich von Andrian^5^, Val Piekarsa^5^, Véronique Giudicelli^19^, Vijay Kuchroo^5^, Woan-Yu Lin^12^, Ziang Zhang^2^

1. NOMIS Center, Salk Institute for Biological Sciences, 2. The University of Chicago, 3. National Institutes of Health, 4. University of California San Diego, 5. Harvard Medical School, 6. Allen Institute for Immunology, 7. Dept of Dermatology and Immunology, University of Pittsburgh, 8. University of Minnesota, 9. University of Washington, 10. UMass Chan Medical School, 11. University of Pennsylvania, 12. Weill Cornell Medicine, 13. The Rockefeller University, 14. Brown University, 15. University of Colorado Anschutz Medical Campus, 16. Swiss Federal Institute of Technology, Lausanne, 17. New York University, 18. La Jolla Institute, 19. IMGT, Univ Montpellier, 20. St. Jude Children’s Research Hospital, 21. Alternative Energies and Atomic Energy Commission, Grenoble, 22. Massachusetts Institute of Technology

## Methods

### Mice

Mice used in the immgenT dataset are described in detail in the immgenT Cosmology manuscript (**Extended Data Table 1**) and on the immgenT website (https://immgen.org/ImmGenT/), including sex, age, and genetic background. With rare exceptions, experiments were performed using C57BL/6 (B6) mice, most of which were sourced from The Jackson Laboratory. Experimental conditions, including infection models, immunization strategies, and tissue processing protocols, are detailed in the immgenT Cosmology manuscript Extended Data Table 1, the GEO GSE297097 dataset, and the immgenT website. Both male and female mice were used.

### ImmgenT dataset: experiments and data processing

The immgenT dataset comprises 66 experiments of single-cell RNA-seq, CITE-seq (128-plex), and paired TCRαβ sequencing (10x Genomics 5′ v2 platform), corresponding to 80 encapsulation runs (“batches”). Each encapsulation is assigned a unique “IGTx” identifier (IGT1-96) used to track datasets. In a minority of cases, two IGT identifiers denote parallel encapsulations of the same biological samples. Individual cells are indexed by a unique IGT.cellID, and samples are tracked using hashtag identifiers (IGT.HT).

A detailed description of data acquisition and processing is provided in the immgenT Cosmology manuscript^12^. Briefly:

#### Logistics

Experiments were conducted across multiple laboratories in the United States. Participating laboratories carried out mouse treatments (e.g., infection or immunization) and tissue processing with their established materials and reagents. The immgenT research assistant (IM) traveled to the site, helped with sample CITE-seq labeling and cell sorting, and performed encapsulation and library preparation. Each experiment typically included ∼10 hashtagged and pooled samples, and a standardized spleen control for batch-effect assessment. Library construction and sequencing were centralized at the Broad Institute.

#### Sample processing, library construction, and sequencing

Flow cytometry-sorted T cells were multiplexed using TotalSeq-C hashtag^67^ antibodies (BioLegend), enabling pooling prior to encapsulation. Cells were stained with a custom 128-antibody TotalSeq-C panel and subjected to joint single-cell RNA, surface protein (CITE-seq), and TCR sequencing using the 10x Genomics 5′ v2 platform. Libraries were sequenced on an Illumina NovaSeq.

*Data processing and quality controls*. Gene expression, protein, hashtag, and TCR count matrices were generated using Cell Ranger, and hashtag-based demultiplexing was performed in Seurat^68,69^. Cells were filtered based on RNA and protein quality-control metrics detailed in the immgenT Cosmology manuscript - Extended Data Note 1.

#### Data integration

Datasets were integrated using totalVI^30,70^, with the 10x Genomics lane (IGT) specified as a batch covariate. The model was trained on all detected genes and proteins using a 30-dimensional latent space (default parameters).

#### Dimensionality reduction

Dimensionality reduction was performed using Minimum Distortion Embedding (MDE) implemented in the pyMDE^71^ library via the pymde.preserve_neighbors() function with default parameters. In contrast to UMAP, which is stochastic and graph-based, pyMDE preserves both local neighborhood structure and global geometry with minimal distortion. After computing the embedding on the full immgenT dataset (IGT1-96), coordinates were reused to anchor immgenT cells as a reference, enabling projection of new datasets without altering the original embedding. This enabled the construction of shared T cell reference embeddings (both all-T and lineage-specific, including CD4-specific embeddings).

#### Cell clustering and annotation

Clustering was performed in the totalVI latent space using the Louvain algorithm (Seurat FindClusters). Clustering solutions were evaluated across resolutions (0.5-4) and optimized using silhouette scores to balance over- and under-clustering. Clusters were merged or split based on consistency across samples and coherence of RNA and protein expression profiles (see immgenT Cosmology manuscript, Extended Data Note 3). In total, 1,710 of 719,580 cells (0.02%) could not be confidently annotated and were labeled as “unclear” and excluded. Small provisional clusters (<1% of cells) are denoted with a “w” prefix.

CD4 T cells were identified as CD4^+^Foxp3^−^ conventional αβ T cells based on combined RNA and protein expression of canonical markers and TFs, including *Cd3e, Trbc1, Trbc2, Trgc1, Trdc, Cd4, Cd8a, Cd8b1, Foxp3* and *Zbtb16*, as well as surface proteins CD3, TCRβ, THY1.2, CD4, CD8A, CD8B, and TCRγδ. Other TCRαβ^+^ CD4^+^ populations, including Zbtb16^+^ T cells (Tz; iNKT and MAIT cells) and CD4^+^Foxp3^+^ Tregs, formed distinct clusters and were excluded from this definition.

#### CITE-seq analysis

The full panel of 128 antibodies and associated quality metrics are provided in **Extended Data Table 4** and **Extended Data Note 1** of the immgenT Cosmology manuscript^12^. Antibody performance was evaluated using RNA-protein correlation and protein dynamic range, stratifying antibodies into high-, intermediate-, and low-performance groups (n = 61, 41, and 26, respectively). Overall, most of the CITE-seq panel performed robustly, enabling dropout-resistant protein quantification that closely parallels flow cytometry and supports both high-dimensional analyses and conventional gating strategies. For flow cytometry-like plots, protein counts were normalized using Seurat’s LogNormalize method.

#### UMAP and clustering of individual IGT datasets

UMAP and clustering were also performed on individual IGT datasets, generating dataset-specific UMAPs used in the Rosetta2 databrowser and in some immgenT manuscripts. These used the standard Seurat functions: NormalizeData(normalization.method = “LogNormalize”, scale.factor = 1e4) %>% FindVariableFeatures(selection.method = “vst”, nfeatures = 2000) %>% ScaleData(features = VariableFeatures(.)) %>% RunPCA(features = VariableFeatures(.), npcs = 50) %>% FindNeighbors(dims = 1:30) %>% FindClusters(resolution = 1).

### Empirical Bayes Matrix Factorization

#### Input normalization and gene filtering

Prior to factorization, counts were normalized to a shifted-logarithmic scale using Seurat LogNormalize with a scale factor set to the dataset-wide mean library size (the mean total RNA count per cell). Genes were then filtered to exclude classes unlikely to reflect coherent transcriptional programs: T-cell-receptor variable, diversity, joining and constant genes (*Trav, Traj, Trac, Trbv, Trbd, Trbj, Trbc, Trgv, Trgj, Trgc, Trdv, Trdj, Trdc*), mitochondrial genes (*mt*-), ribosomal genes (*Rpl, Rps, Mrpl, Mrps, Rsl*), and predicted genes or pseudogenes (*Gm*-prefixed, -*Rik*, and -*ps* genes), in addition to genes not detected in any cell.

#### Factorization model

We applied semi-non-negative empirical Bayes matrix factorization (EBMF)^24^ to the log-transformed gene-expression matrix to infer gene programs (GPs). Let X ∈ ℝ ^*NxP*^ denote the normalized shifted log-expression matrix, with rows corresponding to cells and columns corresponding to genes. Genes and cells with all-zero expression values were removed before analysis. We modeled **X** using *K* = 20 latent components,

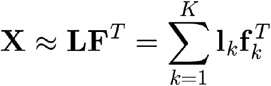

where **L** ∈ ℝ ^*NxK*^ is the cell-by-program loading matrix and X ∈ ℝ ^*PxK*^ is the gene-by-program factor matrix. Here, **l**_*k*_ ∈ ℝ ^*N*^ denotes the activity level of GP *k* across cells, and **f**_*k*_ ∈ ℝ^*P*^ denotes the corresponding pattern of gene regulation across genes.

*Priors*. For each program *k*, the loading vector **l**_*k*_ was assigned an independent scaled point-Exponential prior, defined as a mixture of a point mass at zero and an Exponential distribution with scale parameter 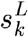. The factor vector **f**_*k*_ was assigned an independent scaled point-Laplace prior, defined as a mixture of a point mass at zero and a Laplace distribution with scale parameter 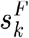. In both cases, the corresponding mixture weights, 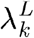 and 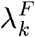, were estimated separately for each program and reflect program-specific sparsity. Residual errors were modeled with gene-specific variances, assumed to be constant across cells within each gene. The model was fitted by empirical Bayes through maximization of the marginal likelihood. These prior choices induce a semi-non-negative factorization, in which loadings are constrained to be nonnegative whereas factors are allowed to take either sign.

#### Model fitting

The computation was done using software *flashier* v.1.0.53 (^72^), where components were first initialized using a greedy iterative strategy and then refined through 200 backfitting iterations; additional implementation details are described in (^24^).

#### Rescaling of gene programs and filtering of outlier cells

For downstream analyses, posterior means of **l**_*k*_ and **f**_*k*_ were used as point estimates. Because **l**_*k*_ and **f**_*k*_ are individually scale-invariant whereas their product 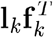 is identifiable, these point estimates were rescaled for interpretability. For each program *k*, **l**_*k*_ was normalized to have maximum value 1, and **f**_*k*_ was normalized to have maximum absolute value 1. Let *d*_*k*_ denote the corresponding scaling constant, and let **D** be the diagonal matrix collecting *d*_*k*_ across programs. The normalized decomposition can then be written as **X** ≈ **LDF**^*T*^ Because this normalization is sensitive to outlier cells with unusually large total loading, we iteratively removed a very small fraction of cells with excessive total program loading before final normalization. Excessive total loading was defined as a total normalized loading greater than 10. In total, only 1,502 cells (0.22%) were removed from the analysis.

#### Reproducibility across datasets

To assess reproducibility, the factorization was re-fit independently within each 10x lane (IGT), and the resulting programs were matched to the reference programs by the cosine similarity of their gene factor scores (over shared genes, after per-program scaling) using a one-to-one Hungarian assignment(RcppHungarian v0.3^73^ ). Batch effects were additionally evaluated on the standardized spleen control included in every experiment. For each program, the proportion of its loading variance explained (pve) by dataset was computed as the one-way ANOVA η^2^ — across all 47,253 spleen-control cells, and compared with the proportion explained by sub-lineage cluster (level 2) over the same cells.

### Characterization of gene programs

#### Highly active cells and genes

Using the rescaled loadings and factor scores (normalized per program to a maximum of 1 and a maximum absolute value of 1, respectively), a cell was defined as highly active in a program when its normalized loading exceeded 0.1, and a gene as highly active when the absolute value of its normalized factor score exceeded 0.25. Each program was summarized by its proportion of highly active cells and its number of highly active genes, and each cell by its number of highly active programs (the programs with normalized loading above 0.1). The number of highly active programs per cell was compared across T-cell lineages and between resting and activated cells, and related to surface CD44 protein levels by Pearson correlation.

#### GP-gene signatures

The signature of each program was defined from its normalized factor scores, taking genes with a score above 0.1 as positive (up-regulated) signature genes and genes with a score below −0.1 as negative (down-regulated) signature genes, ranked within each direction by the magnitude of their score (**Extended Data Tables 1 and 2**).

### Gene programs discriminating lineage, tissue, and cluster identities

For each gene program and each cell category, we quantified how well the program’s cell loading discriminated that category by a one-vs-rest receiver-operating-characteristic analysis (pROC v1.19.0.1^74^), treating cells in the category as positives and all other cells as negatives and using the program loading as the predictor; the area under the curve (AUC) summarizes the discrimination. For threshold-based analyses, the optimal loading cutoff was taken as the point on the ROC curve closest to the top-left corner. AUCs were computed for three category types — cell lineage (level 1), sub-lineage cluster (level 2), and tissue of origin — and, throughout, on healthy, non-thymocyte cells.

The complete one-vs-rest AUC matrices for lineage, tissue, and cluster are provided in **Extended Data Tables 3, 5 and 6**. A program was taken to positively predict a category when its AUC exceeded 0.8 and its optimal loading threshold was at least the program’s median loading, so that high, rather than low, loading drove the prediction; these per-program calls are summarized in **Extended Data Table 1**. Lineage-, tissue-, and cluster-specific programs were identified by restricting each program to the categories in which it was up-regulated (category mean loading above the overall mean) and taking its maximum AUC across those categories (excluding categories below a size floor of 100 cells for tissues and clusters), and programs with a maximum AUC above 0.9 were taken as category-specific.

### Gene programs during T-cell activation

#### Differential loading and standardized effect sizes

For each GP, the activation-associated change was quantified as the difference in mean cell loading between activated and resting cells, computed separately within CD4 and within CD8. To place GPs on a common scale, we also computed a standardized mean difference by dividing this change by a pooled standard deviation of the loading estimated across all resting and activated CD4 and CD8 cells. Programs were then classified by the concordance of their CD4 and CD8 responses — induced predominantly in one lineage when the two changes differed by more than three-fold, or in both lineages otherwise — subject to a minimum standardized effect size of 0.15. These per-program statistics are reported in **Extended Data Table 4**. Where fold-changes in mean loading are reported, they were computed as log2 fold-changes with a pseudocount of 10^−10^.

#### Gene-set enrichment analysis

Program gene signatures were tested for enrichment of the MSigDB^64^ C7 immunologic signature collection (mouse orthologs) using the perform_gsea function of the pathways R package, with each program’s gene factor scores as the ranking statistic; normalized enrichment scores and p-values were computed per program and gene set, and p-values were converted to q-values (qvalue).

### Surface protein expression associated with GPs

#### Protein-program projection

To assign surface-protein signatures to the transcriptionally defined programs, the CITE-seq data were projected onto the fixed scRNA-derived program loadings. Antibody-derived-tag (ADT) counts were log-normalized (with a scale factor equal to the rounded mean ADT count per cell) to form a cell-by-protein matrix, which was modelled as the product of the fixed cell loadings and a program-by-protein factor score matrix. The protein scores were estimated by empirical Bayes matrix factorization (flashier v1.0.53^31^) with the cell loadings held fixed at their scRNA values, using a point-Laplace prior on the protein scores and protein-specific residual variances, initialized by ordinary least squares and backfit for 200 iterations. Analyses were restricted to CITE-seq-measured cells and to the 47 quality-passing antibodies of the 128-antibody panel (antibody quality control is detailed in the immgenT Cosmology manuscript^12^). The protein score matrix was normalized per program to a maximum absolute value of 1.

#### Protein-gated embedding plots

For visualization, the protein-gated cells and an equally sized set of top-loading cells were displayed on the shared MDE embedding and colored by local point density; thymocyte, proliferating and miniverse cells were excluded from these comparisons.

#### KLRG1 and CD69 associations

Individual proteins were related to program activity: for KLRG1, each program’s mean loading was compared between KLRG1-positive and KLRG1-negative cells within CD8, CD4 and Treg cells; for CD69, the Spearman correlation between program loading and CD69 protein level was computed.

### Residual Quantized Variational Inference (RQVI)

#### Overview of the algorithm

To uncover interpretable gene programs from single-cell RNA-sequencing data, we developed Residual Quantized Variational Inference (RQVI), a generative model built on combination of scVI^59^ and RQ-VAE^75^. Continuous latent embeddings like those of scVI are suitable for prediction but give every cell its own private representation, which is difficult to interpret biologically. RQVI augments this continuous representation with a discrete bottleneck: each cell is expressed as a sum of shared “gene-program” vectors drawn from a learned dictionary, so the same programs are reused across all cells. Each program is therefore encouraged to capture a transcriptional module that recurs in the data and can be characterized by the genes it most strongly affects.

#### Gene-program dictionary

Biological variation in single-cell data is naturally hierarchical, ranging from broader to subtle context-specific states. To respect this structure, RQVI represents each cell through *D* successive refinements of its encoder embedding. Let zi ∈ ℝ^*H*^ be the embedding of cell *i*, let *C* = {*e*_1_, …, *e*_*K*_} be a shared dictionary of *K* code vectors, and initialize the residual as 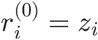 . At refinement depth *d*, RQVI selects the code vector closest to the current residual and then subtracts that vector to obtain the residual for the next depth:

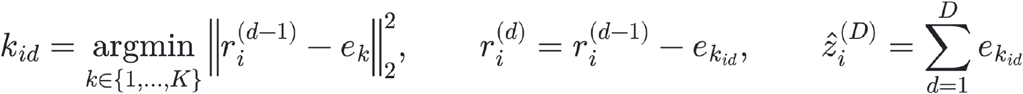

At the first depth, the residual is the complete encoder embedding, so the selected code typically captures the dominant signal. Once this contribution has been removed, the next code is selected against what remains; successive depths therefore explain only the variation missed by earlier ones. Summing the *D* selected vectors gives 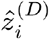, a coarse-to-fine representation built from a small vocabulary of reusable components.

Following RQ-VAE, the same codebook *C* is used at every depth. Sharing the codebook requires only a single codebook-size parameter *K* and makes every code available at every refinement step, maximizing the utility of the learned vocabulary (Lee et al., 2022). Because the residual being approximated changes with depth, depth is part of the meaning of a code assignment: a code selected at the first depth represents a coarse component of cell state, whereas the same code selected at a later depth represents a refinement conditional on the preceding selections. We therefore define a gene program by its depth–code pair (*d, k*). With *K* = 64 possible codes at each of *D* = 4 depths, RQVI yields 64 programs per depth and *D*×*K* = 256 gene programs in total.

#### Dictionary learning and training schedule

Dictionary entries are updated as a running average of the embeddings each entry is responsible for, which lets the programs track the data without competing with the gradients flowing through the encoder. Entries that go unused are reseeded from a noisy copy of an active embedding to prevent collapse onto a few dominant programs, a known failure mode of discrete bottlenecks. Training also proceeds progressively, allowing only one program per cell at the start and activating additional programs at fixed milestones, because enabling all depths at once lets the later programs absorb structure that the earlier ones should learn. The objective then combines (i) reconstruction of the gene-expression counts, (ii) the standard scVI regularisation terms on the cell embedding and per-batch library size, and (iii) a penalty pulling the encoder embedding toward its assigned dictionary entries.

#### Implementation

RQVI was trained with a 256-dimensional cell embedding (*H* = 256), hidden layer of size 512, K = 64 codebook size program vectors, D = 4 refinement steps, dropout 0.1, batch size 256, and 30 epochs, using the Adam optimiser through the scvi-tools training loop on a 90/10 training/validation split. The choice of K = 64 and D = 4 balances expressive capacity against interpretability: it provides enough programs to span the heterogeneity of our data while still allowing every program to be inspected manually.

#### Cell-level program loadings

Residual quantization used hard nearest-code assignments during model fitting to avoid overfitting. For downstream analysis, we calculated soft assignments from the negative squared distances to the 64 code vectors at each depth using a softmax with temperature 1. The four sets of 64 depth-specific loadings were concatenated to form the cell-by-program loading matrix.

#### Linking gene programs to genes

Gene effects were estimated from the trained decoder using a residual-delta calculation. For each depth, a reference latent vector was constructed from the mean codebook contributions at the preceding depths. For a depth-code program (*d, k*), decoder-predicted expression was evaluated before and after adding code vector *e*_*k*_ to this reference. Their difference on the log(1 + *x*) scale defines the signed gene-effect vector. Because the reference represents the components already selected at earlier depths, the resulting gene effect describes the additional transcriptional contribution of the code at its specified depth.

#### RQVI scalability benchmark

RQVI was trained on subsampled datasets of approximately 10k, 50k, 100k, 200k, 400k, and 633k cells (the full ImmGen T dataset), with 3 replicates per size. Model architecture and fitting parameters were held fixed. Wall-clock time was measured for model training. Mean training time increased approximately linearly with the number of cells (R^2^ ≈ 0.99). Training on full datasets required approximately 35.5 min on a single NVIDIA RTX A4000 GPU.

#### RQVI GP sparsity metrics

For each of the 256 RQVI GPs (seed 0), we computed: (i) the proportion of active cells (fraction of cells with loading > 0.01); (ii) the number of active genes, defined as genes with |W_scaled| > 0.45 after global min-max normalization of the gene-effect matrix W to [−1, 1]; and (iii) a per-GP percentage of variance explained (PVE), computed as the product of the squared norms of each GP’s cell-loading and gene-effect vectors, normalized to sum to 1 across all 256 GPs. PVE was displayed on a log scale.

#### Comparing EBMF and RQVI factors

EBMF programs were benchmarked against independent sets of gene programs derived by RQVI, which were generated across ten independent random seeds (256 programs per seed; 2,560 candidate factors in total). For the comparison, each program’s mean loading profile across level-2 clusters was z-scored, EBMF and RQVI programs were compared by the signed Pearson correlation of these profiles, and the 200 EBMF programs were matched one-to-one to the pooled RQVI factors by optimal (linear-sum) assignment maximizing correlation (SciPy v1.15.3).

### Plotting

#### Software and pseudo-counts

Plots were generated in R using ggplot2^76^, S-Plus, Seurat^69^, or the ZemmourLib R package (https://github.com/dzemmour/ZemmourLib, v0.1.3). Heatmaps were generated using pheatmap (v1.0.13) or Morpheus (Broad Institute). Where fold-changes in mean program loading are shown, they were computed as log2 fold-changes after adding a pseudo-count of 10^−10^ so that zero values remain defined.

#### Structure plots

Structure (admixture) plots of per-cell program loadings were drawn with fastTopics (v0.7.25) on a subsample of ∼100,000 cells.

#### Alluvial plots

Alluvial diagrams were drawn with ggalluvial (v0.12.6), using program-positive cells subsampled to 300 per program with negligible flows omitted, so that ribbon widths reflect relative rather than absolute cell numbers.

Panel-specific plotting details are available in the project code repository.

## Code availability

Code is available at the following repository https://github.com/immgen/immgenT_Project/tree/main/GP_paper

## Data availability

All raw and processed sequencing data generated in this study are available through the Gene Expression Omnibus (GEO) under accession GSE297097.

## immgenT resource: public data browsers

Beyond the data and analyses reported in this and accompanying articles, immgenT’s goal as a public resource is served by a set of data browsers which enable multi-faceted exploration of the data. They are accessible through the immgenT portal (https://www.immgen.org/ImmGenT/) and include:

**Rosetta2 (https://rosetta.immgen.org/)** is an interactive single-cell viewer that connects RNA-seq and protein (CITE-seq) data. Cell populations can be visualized on parallel RNA and protein UMAPs, coloring reciprocally RNA- or protein-based clusters. The expression of individual genes, or of user defined gene signatures can be mapped on these representations. A cell gating interface modeled after flow cytometry serially generates 2D CITE-seq plots, on which cells can be “gated”, for projection onto UMAPs or computation of differential expression. Rosetta2 returns differentially expressed genes and proteins (as heatmaps or tables). Rosetta2 can be used to explore individual datasets or, or integrated cells grouped by lineage.

**TCR Browser** (https://rstats.immgen.org/tcrbrowser/) is a searchable interface for TCR sequences across the immgenT data.

TCR repertoires can be browsed within immgenT datasets, searched for TCRs that use specific V, J or CDR3 elements, or for similarity to a user provided TCR.

**T-RBI portal** (Reference Based Integration) (https://rstats.immgen.org/immgenT_integration_analysis/) allows AI-driven mapping of external single-cel RNAseq datasets into the immgenT framework. Users upload their own data and metadata files (standard sc formats), which are then mapped and annotated within the immgenT framework. After an overnight compute, the tool returns annotation tables with integration statistics and annotation of every cell in the immgenT cluster framework, and x/y coordinates in the reference MDE space.

**Skyline browser**

(https://rstats.immgen.org/Skyline/skyline.html?datagroup=immgenT%20Pseudobulk&) represents, as a classic bar histogram, a pseudo-bulked overview of expression profiles of a given gene across all or selected immgenT cell clusters

## Extended Data Legends

**Extended Data Figure 1. GP reproducibility and batch-effect evaluation**.

**(a)** Cumulative number of GPs, out of the 200 identified in the full immgenT solution, reproduced in at least one dataset (IGT). For each dataset, an EBMF factorization was computed and its factors were matched to the 200 immgenT GPs by Hungarian assignment using the cosine similarity of gene-score vectors. A GP was considered reproduced in a dataset when this cosine similarity exceeded the threshold indicated for each curve.

**(b)** Distribution of GP reproducibility across datasets. Curves show the number of GPs reproduced in at least that many datasets, for cosine-similarity thresholds ranging from 0.2 to 0.8.

**(c-e)** Batch-effect evaluation using spleen standards (T cells in 6-8 week old mice, unchallenged).

**(c)** For each GP, the proportion of loading variance explained by datasets (IGT) (x-axis) versus the proportion explained by T cell clusters (level-2 cluster, y-axis), computed over all spleen-standard cells, identifies batch-effect in GPs.

**(d)** In some instances, GP captures specific batches, such as GP9, specific to IGT13-14. Boxplots showing the distribution of GP9 activity in the spleen-standard cells across datasets.

**(e)** In other instances, like GP1, GPs capture continuous batch effects such as sequencing depth. Mean RNA count per dataset (x-axis) against mean GP1 loading per dataset (y-axis) in the spleen-standard cells.

**(f)** For each GP, the number of active genes versus the fraction of active cells (log-scaled).

**Extended Data Figure 2. Lineage and cluster-associated GPs**.

**(a)** Box plots showing the distribution of GP30 activity across Tz subsets—iNKT, MAIT, and other Tz cells (that is, Tz cells that are neither iNKT nor MAIT)—and all other T cells.

**(a)** Box plots showing GP58 activity in resting and activated CD8+ T cells compared with all other T cells.

**(c)** Box plots of CD8A and CD8B protein expression (CITE-seq, log-normalized counts) in resting and activated CD8+ T cells.

**(d)** Row-centered mean GP activity across clusters. Mean GP activity computed in samples at baseline.

**Extended Data Figure 3. Activation-associated GPs**.

**(a)** Definition of resting and activated CD4+ and CD8+ T cells. Activated CD44^+^CD62L^−^ (top) and resting CD62L^+^CD44^−^ (bottom) cells are highlighted in each lineage-specific MDE. Gated cells are shown in the adjacent CD62L versus CD44 protein-expression plot (CITE-seq, log-normalized counts).

**(b)** Bar plot showing broad GP26 activity in activated CD4+ and CD8+ T cells across organs (percent of cells with GP26 activity above 0.1 per tissue)

**(c)** Gene set enrichment analysis (GSEA) dot plot relating activation-associated GPs to curated immunologic signature gene sets. Color denotes −log10 adjusted *P* value, and dot size denotes the normalized enrichment score (NES).

**(d)** Bar plot showing the percentage of GP79-active cells among all activated CD4+ T cells across immunological challenges.

**(e)** Heatmap showing single-cell GP activity in activated CD4+ and CD8+ T cells (log2 fold change in GP activity relative to the resting baseline of the same lineage). Cells are grouped by cluster.

**(f)** Bar plot showing the percentage of GP57-active cells among activated CD8+ and CD4+ T cells in cancer versus all other conditions.

**Extended Data Figure 4. Protein expression across GPs**.

Heatmap of scaled protein scores for each GP. We focused on the 47 proteins that performed best in the immgenT CITE-seq dataset.

**Extended Data Figure 5. Examples of surface proteins associations with GP**.

**(a,b)** Mean activity of the ten CD69-associated GPs (a) across tissues and (b) across lineages. **(c-f)** Examples of gating strategies used to identify GP-active cells for (c) GP29 (CD8aa gdT or ab T cell specific), (d) GP58 (CD8-specific), (e) GP22 (DN-specific), and (f) GP68 (Treg-specific).

**Extended Data Figure 6. RQVI scalability and representative matches to lineage-associated EBMF GPs**.

**(a)** RQVI training time as a function of the number of input cells. The dashed line indicates the linear fit.

**(b)** Scatter plot summarizing sparsity and variance explained for RQVI GPs. Each point represents one RQVI GP. The x axis shows the proportion of active cells, the y axis shows the number of active genes, and color indicates the percentage of variance explained.

**(c-f)** All-T UMAP plots showing RQVI GP activity for representative matches to lineage-associated EBMF factors: (c) F22 matched to RQVI GP96 from seed 6, (d) F30 matched to RQVI GP112 from seed 5, (e) F58 matched to RQVI GP110 from seed 1, and (f) F68 matched to RQVI GP76 from seed 3. Pearson correlation coefficients for the matched pairs are shown above each plot. Color indicates GP activity from low to high.

**(g-j)** Gene-effect plots for the matched RQVI GPs shown in (c-f): (g) GP96, (h) GP112, (i) GP110, and (j) GP76. Each point represents a gene. The x axis shows the RQVI gene effect, and the y axis shows mean expression. Selected high-effect genes are labeled.

## Extended Data Tables

**Extended Data Table 1 - GP summaries and annotations**

Characteristics of the 200 GPs: annotations, proportion of highly active cells (scaled loading > 0.1), and number of highly active genes (|score| > 0.25).

**Extended Data Table 2 - Gene signature per GP**

Signature genes of all 200 GPs, defined as genes with score > 0.1 (up) and < -0.1 (down) and capped at the 100 highest-scoring genes per GP and direction.

**Extended Data Table 3 - GP association with lineages**

AUC score separating cells of each lineage from all others using GP loadings (computed in cells at baseline).

**Extended Data Table 4 - GP activity during activation**

Change in GP loading with T cell activation, computed in CD4 and CD8 cells separately: the mean loading in activated minus resting cells; d_CD4 and d_CD8, the corresponding standardized differences, each divided by the s.d. of the GP’s loading across all activated and resting CD4 and CD8 cells; and Ratio_CD8_CD4, the ratio of the CD8 to the CD4 mean loading change.

**Extended Data Table 5 - GP association with tissues**

AUC score separating cells of each tissue from all others using GP loadings (computed in tissues at baseline).

**Extended Data Table 6 - GP association with clusters**

AUC score separating cells of each cluster from all others using GP loadings (computed in cells at baseline).

