## Extended Data Figure for "A Gene-Program Architecture of Mouse T cells"

### Extended Data Figure 1

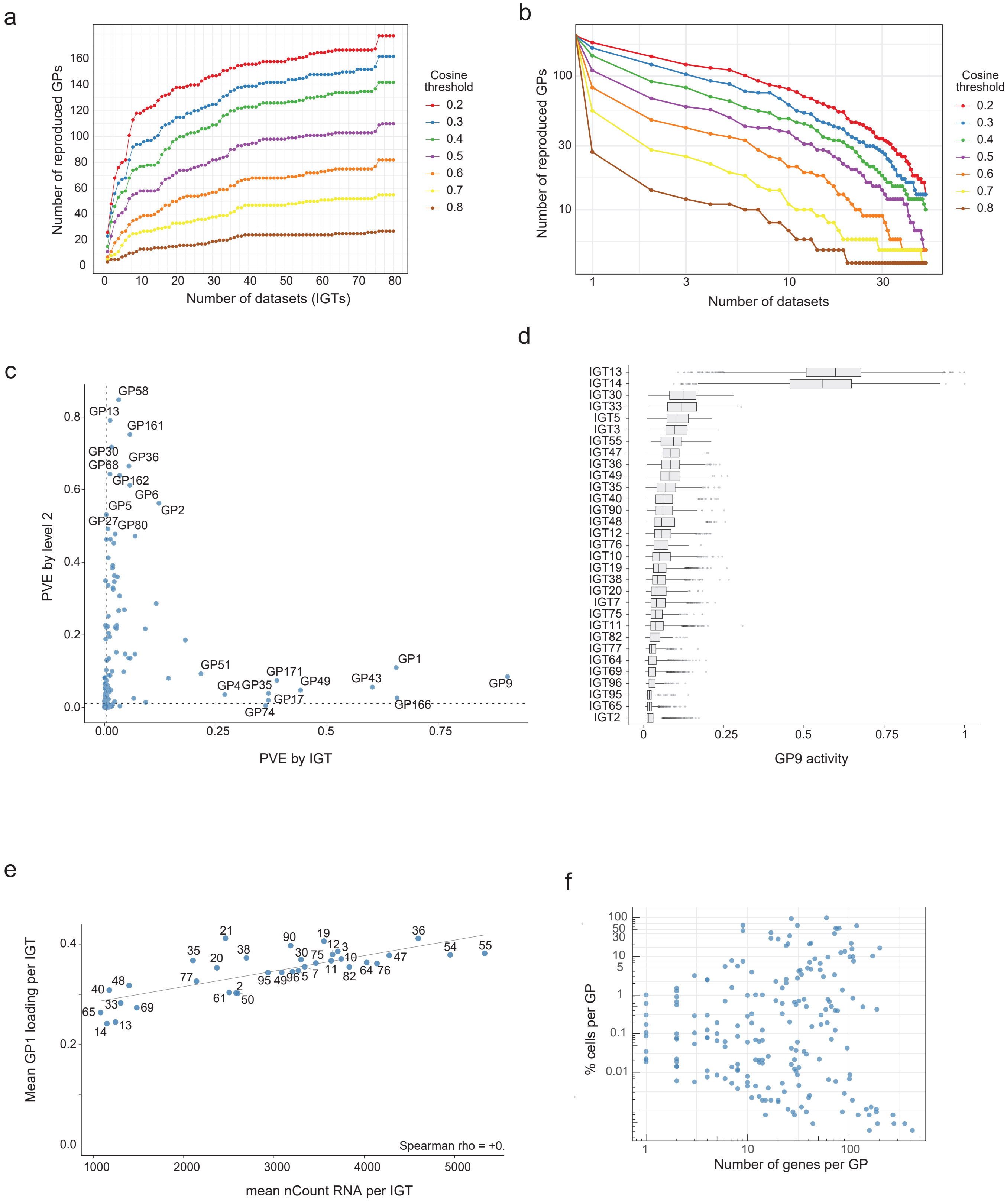

### Extended Data Figure 2

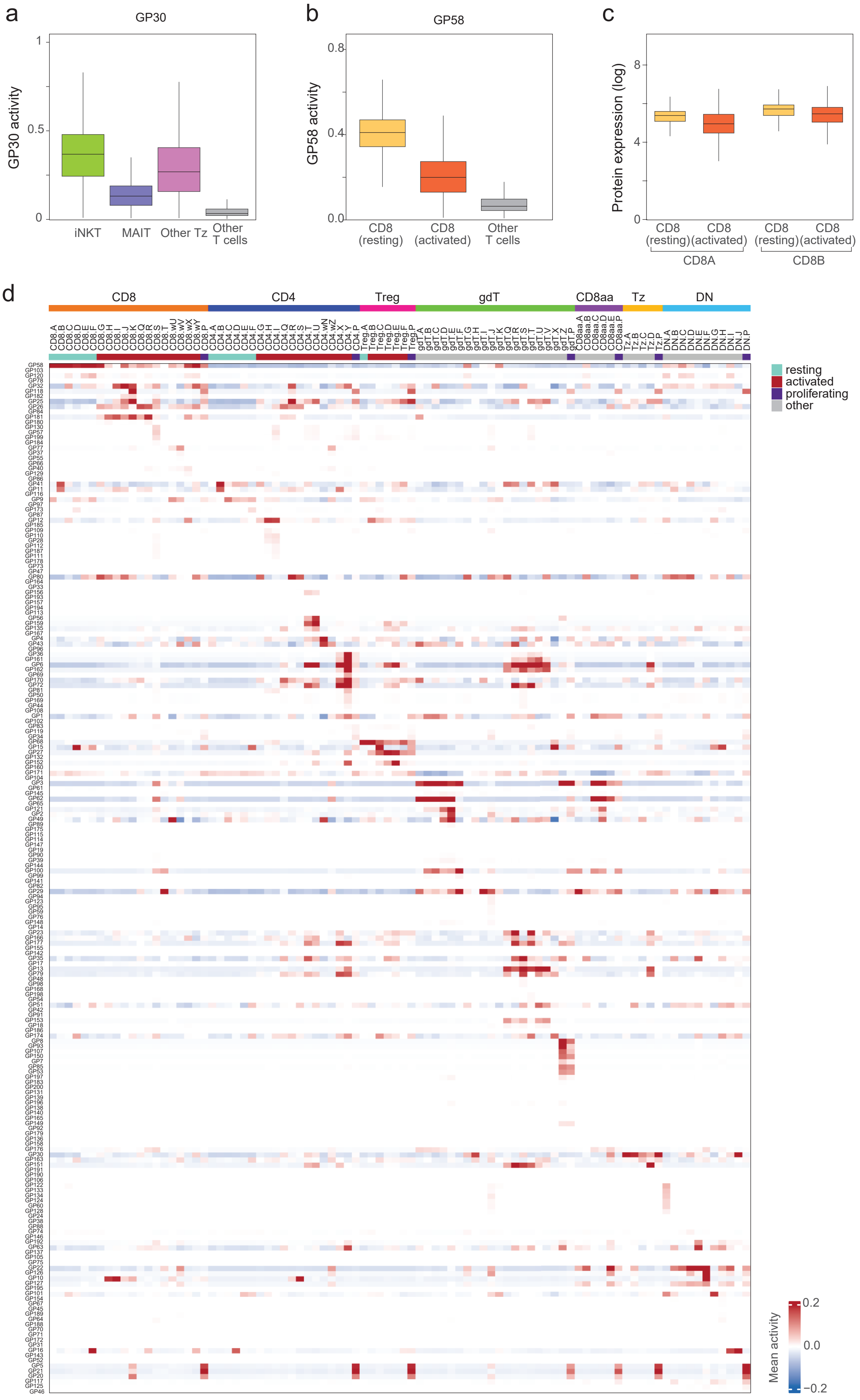

Extended Data Figure 3

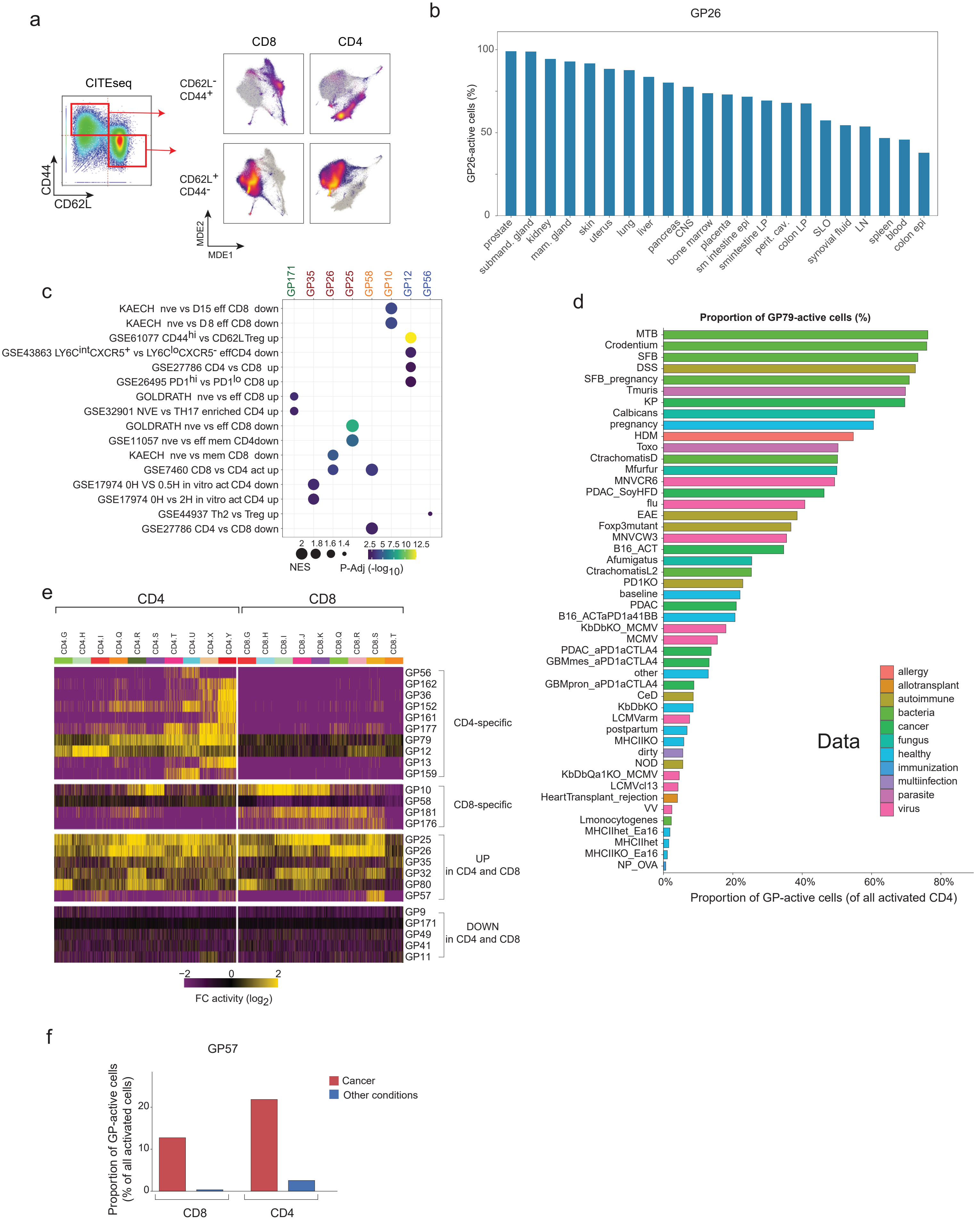

### Extended Data Figure 4

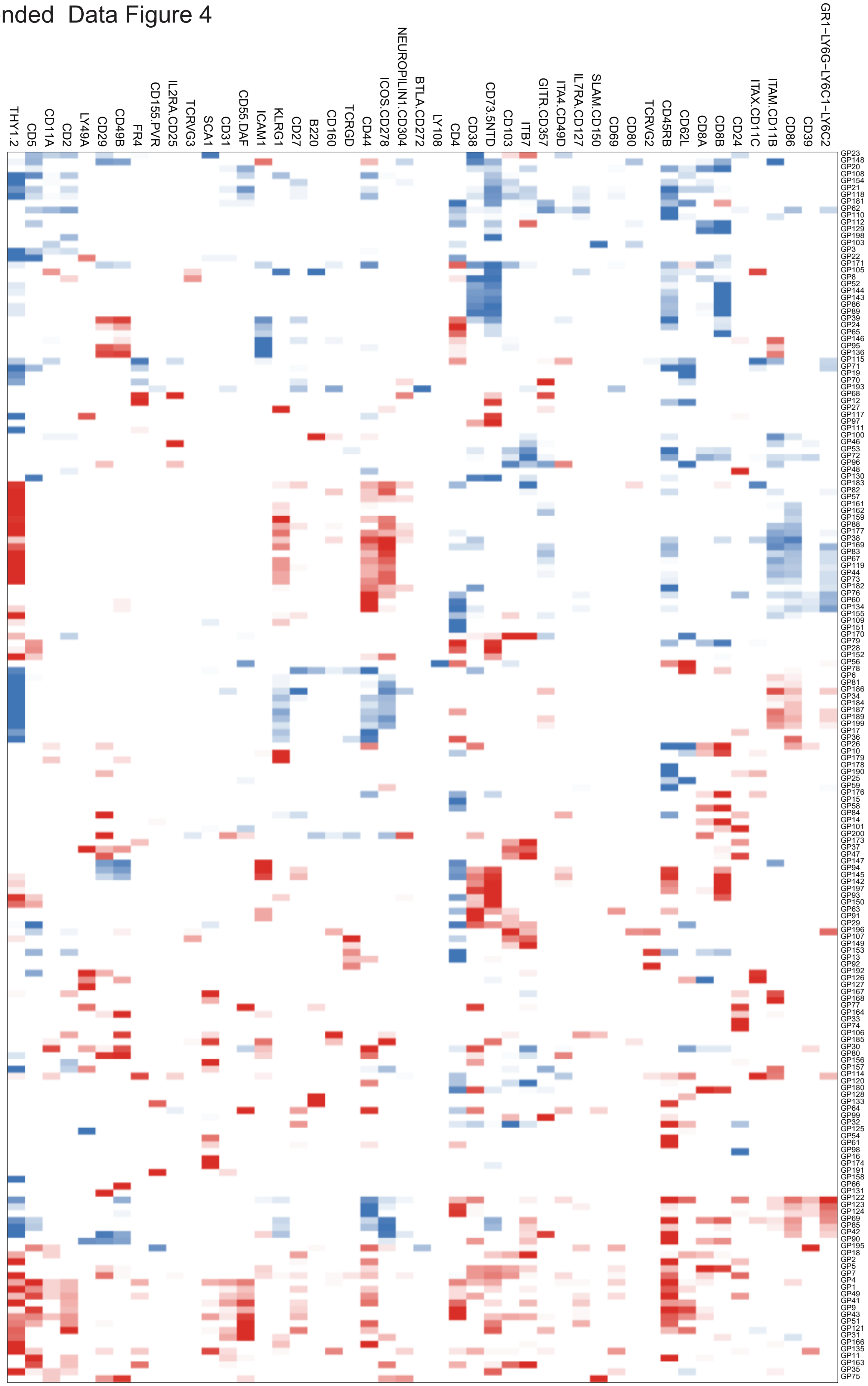

### Extended Data Figure 5

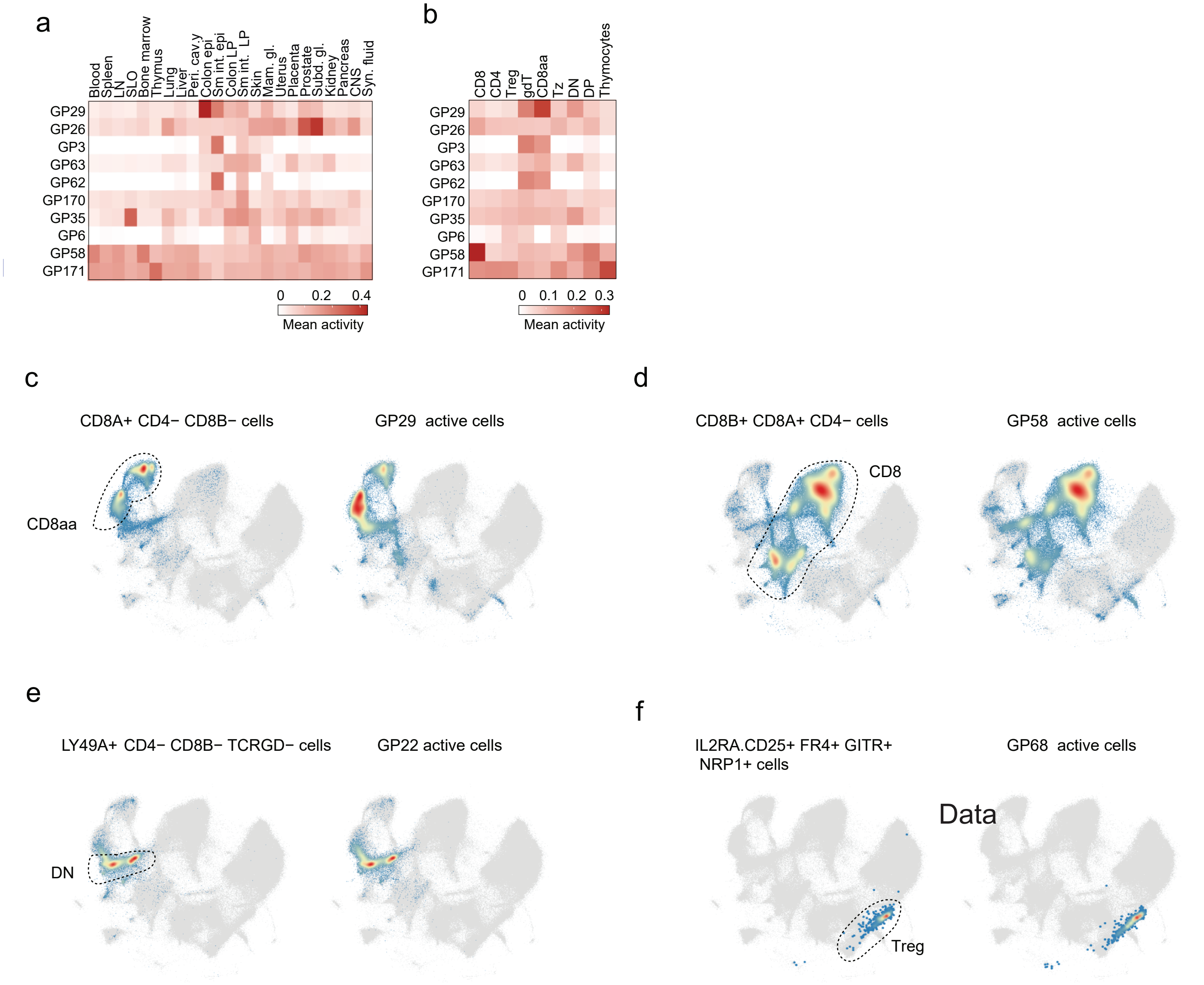

### Extended Data Figure 6

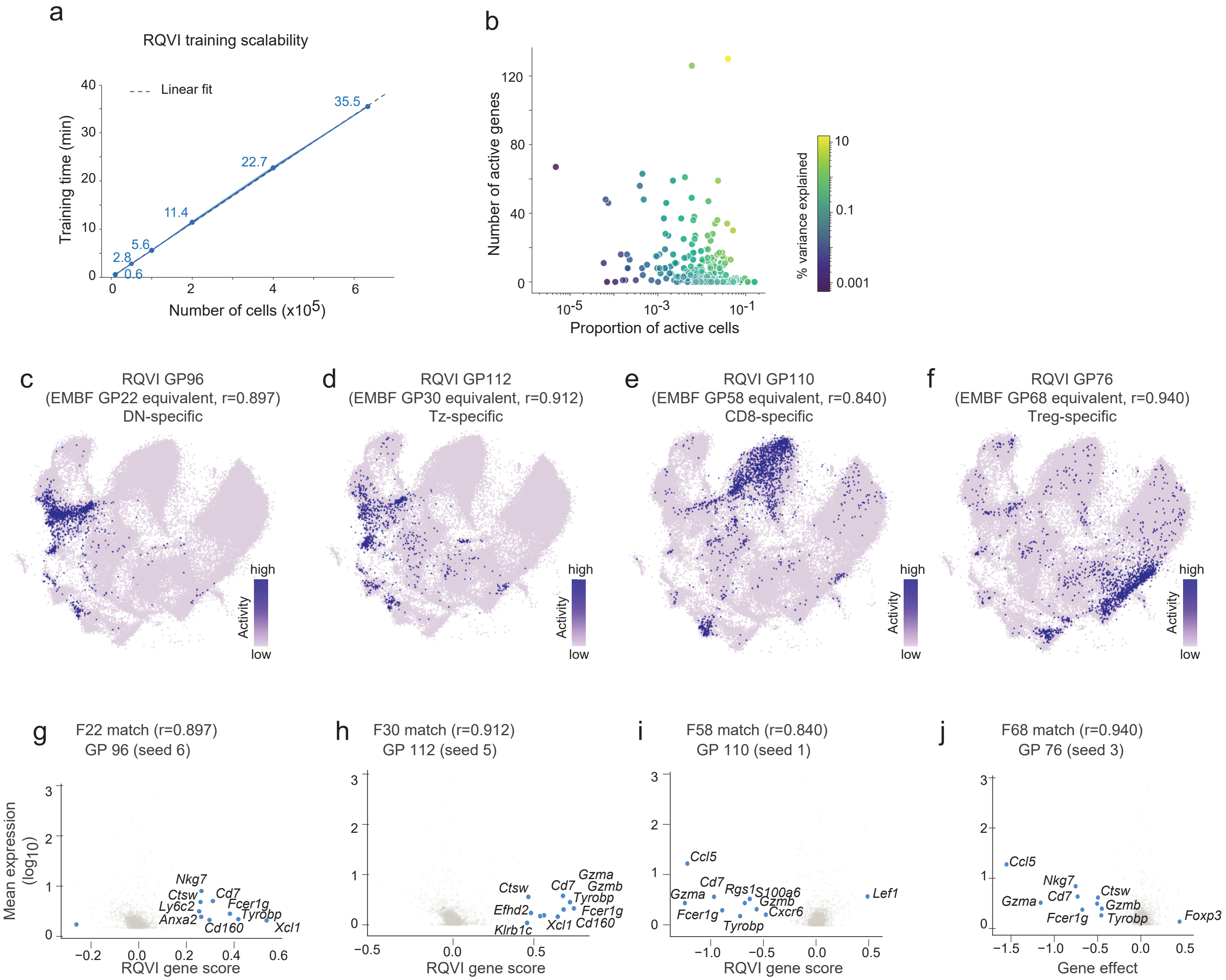
